# Asymmetric control of food intake by left and right vagal sensory neurons

**DOI:** 10.1101/2023.05.08.539627

**Authors:** Alan Moreira de Araujo, Isadora Braga, Gabriel Leme, Arashdeep Singh, Molly McDougle, Justin Smith, Macarena Vergara, Mingxing Yang, M Lin, H Khoshbouei, Eric Krause, Andre G de Oliveira, Guillaume de Lartigue

## Abstract

We investigated the lateralization of gut-innervating vagal sensory neurons and their roles in feeding behavior. Using genetic, anatomical, and behavioral analyses, we discovered a subset of highly lateralized vagal sensory neurons with distinct sensory responses to intestinal stimuli. Our results demonstrated that left vagal sensory neurons (LNG) are crucial for distension-induced satiety, while right vagal sensory neurons (RNG) mediate preference for nutritive foods. Furthermore, these lateralized neurons engage different central circuits, with LNG neurons recruiting brain regions associated with energy balance and RNG neurons activating areas related to salience, memory, and reward. Altogether, our findings unveil the diverse roles of asymmetrical gut-vagal-brain circuits in feeding behavior, offering new insights for potential therapeutic interventions targeting vagal nerve stimulation in metabolic and neuropsychiatric diseases.

**One Sentence Summary:** Lateralized gut-brain circuits respond to different sensory modalities and control distinct feeding behaviors.

## Introduction

The ability to efficiently satisfy metabolic requirements while maximizing time devoted to other important behaviors is critical for survival. To achieve this, substantial cognitive effort needs to be dedicated to assessing what to eat, where to find it, and how much to consume. The vagus nerve provides a primary neural mechanism for sensing and integrating post-ingestive signals that influence acute and long-term decisions related to feeding behaviors (*1, 2*). While significant advances have been made in phenotyping the morphology (*3–6*) and molecular profile (*7–10*) of vagal sensory neurons of the nodose ganglia (NG), the fundamental organization for integrating and processing the variety of gut information that produces competing behaviors remains unclear. Based on evidence that asymmetry of neural circuits can enhance integration of multiple simultaneous sensory cues (*11, 12*), we propose the hypothesis that lateralized processing of gastrointestinal information by nodose ganglia could provide a competitive advantage to optimally guide feeding decisions by engaging different feeding-related behaviors and circuits. Our research utilizes molecular and genetic approaches to record and manipulate the activity of vagal sensory neurons to identify separate roles for the left and right vagus nerve in coordinating feeding behavior. Our findings offer new insights into how the brain processes gut information and provide a better understanding of the mechanisms involved in the regulation of feeding behavior.

## RESULTS

### Profiling of NG neurons predicts vagal asymmetry

The molecular identity of anatomically and functionally defined vagal sensory neurons are now well cataloged (*7–10*). However, the extent to which gene expression patterns differ between the LNG and RNG neurons has been largely overlooked. We used publicly available targeted scRNAseq data (*13*) to reveal the molecular constituents that may underlie differences between left and right NG neurons. Vagal sensory neuron clusters were isolated using the previously defined *Phox2b* gene marker, excluding other cell types (Fig. S1A and S1B).

Cocaine- and Amphetamine-Regulated Transcript (CART) is a neuropeptide widely expressed in the NG (*14*). CART expression increases in response to nutrient availability with the change being more pronounced in the RNG (*15*). RNAseq data confirms that the *Cartpt* gene is ubiquitously expressed gut-innervating vagal sensory neurons (*16*), making it an attractive genetic target for studying asymmetry of vagal mediated gut-brain signaling.

We confirmed that *Cartpt* neurons are broadly expressed in NG clusters (Figure 1A) and are equally represented in left and right NG (Fig. S2A-B). Analysis of NG*^Cart^* neurons resulted in striking gene expression pattern segregation between the left and right NG. LNG*^Cart^*neurons preferentially co-expressed oxytocin receptor, a molecular marker of mechanosensory vagal neurons that form intestinal IGLE (*9*). Furthermore, these neurons highly co-express the mechanically sensitive channels *Piezo1* and *Piezo2* (Figure 1B). In contrast, RNG*^Cart^* neurons preferentially co-expressed *Vip* and *Gpr65*, markers of chemosensory vagal neurons that innervate small intestine villi (*9*). Notably, while asymmetric gene expression is preferentially enriched in NG*^Cart^* neurons, this also translates more generally to all *Phox2b* expressing vagal sensory neurons (Figure S1C). These separate gene expression patterns predict anatomical and functional asymmetry of the vagus nerve.

**Figure 1.**
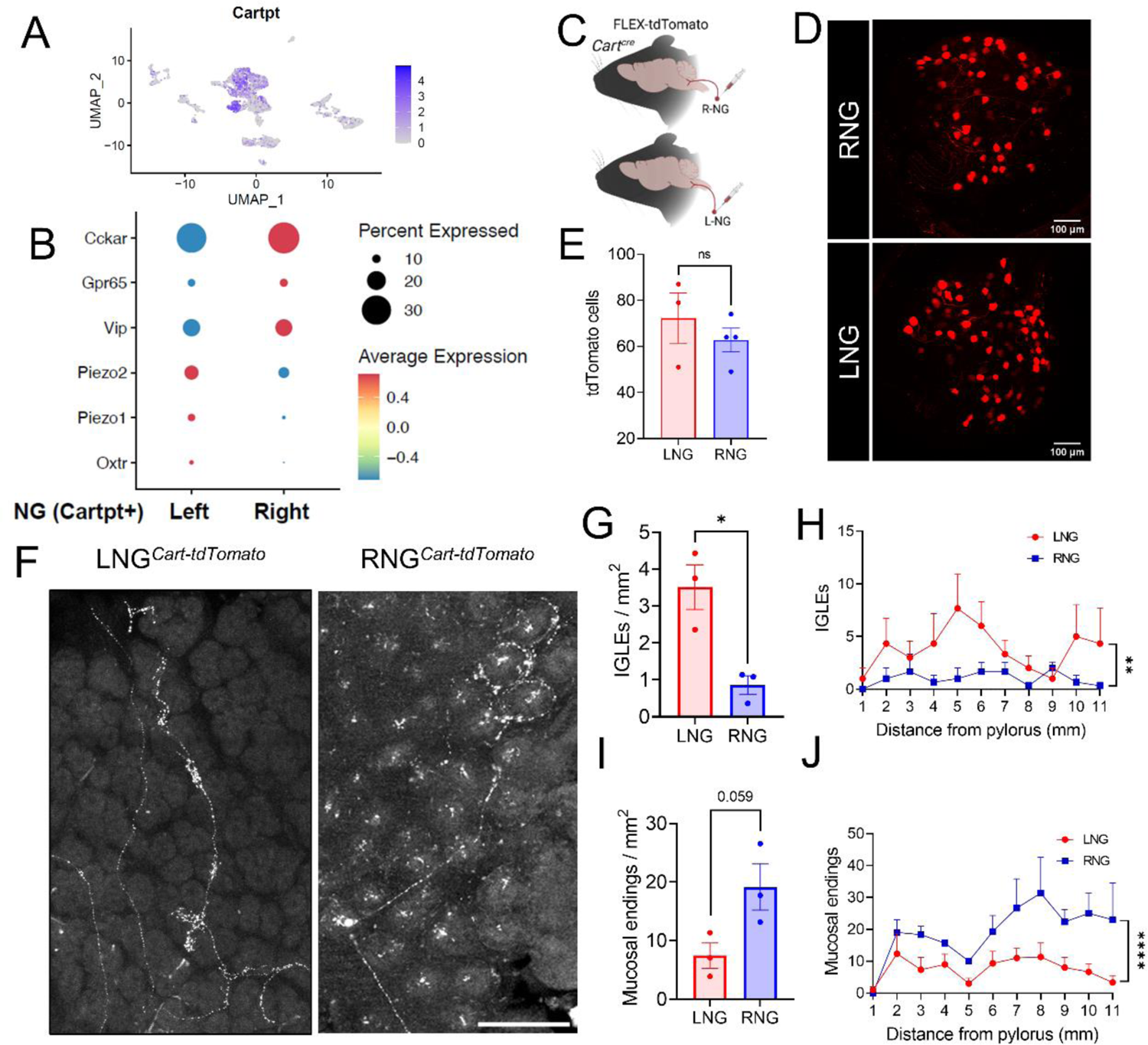
LNG^Cart^ and RNG^Cart^ Asymmetrically Innervate the Duodenum. (A) subpopulation of NG neurons that express the *Cartpt* gene within the *Phox2b* cluster. (B) Dot plot showing expression of selected genes involved in mechanosensing and nutrient sensing in NG*^Cart^* neurons. (C) Unilateral DIO tdTomato injection (n=3-4). (D) tdTomato detected in LNG and RNG. (E) Quantification of (C). (F) Whole mount imaging of the duodenum showing vagal terminals from LNG*^Cart^* or RNG*^Cart^* fibers. (G-J) Quantification of IGLEs and mucosal endings in the duodenum (n=3, unpaired Student’s t test and two-way ANOVA). Data are expressed as mean ± SEM; ns, p > 0.05; t tests and post hoc comparisons, *p < 0.05. Scale bars 100 µm.

To study CART expressing NG neurons, we validated a newly developed *Cart-Cre* mice as a viable genetic tool for targeting vagal sensory neurons that express the *Cartpt* gene (Fig. S2C-D). Previous studies have identified a number of different vagal terminal endings in the intestine and shown that these terminal morphologies predict their function (*3–6, 17*). Intra-ganglionic laminar endings (IGLEs) and intramuscular arrays are specialized sensory terminals that detect mechanical stimuli (*18*) and are present in higher numbers in the proximal intestine (*4, 5*). In contrast, mucosal terminals found surrounding the intestinal crypts or as free terminals embedded within the lamina propria of intestinal villi (*3, 6, 9, 19*) sense chemical stimuli (e.g., nutrients) (*20*). To assess where vagal CART neurons project in the gastrointestinal tract, we used a viral-guided mapping strategy. Cre-dependent AAV encoding fluorescent reporter (_AAV_**_PhP.S_**-DIO-tdTomato) was injected unilaterally into NG of *Cart-Cre* mice (Fig. 1C), and fibers were visualized using 3D imaging of cleared whole mounts from proximal intestine. The LNG and RNG exhibited comparable number of tdTomato+ neurons (Fig. 1D,E), and the LNG*^Cart^* and RNG*^Cart^*terminals on the dorsal vagal complex were mostly lateralized. Nevertheless, fibers were seen crossing over into the AP and contralateral NTS at certain points along the caudal-rostral length (Fig. S2E). In the intestines, vagal fibers penetrated the duodenum through the mesenteric attachment and coursed toward the antimesenteric pole (*3*), and NG*^Cart^* fibers densely innervated both the muscular layer and intestinal mucosa (Fig. 1F). The quantification of duodenal sensory terminals in the first 11 mm of the mouse intestine revealed that the LNG*^Cart^*neurons preferentially innervated the muscular layer (Fig. 1F) and accounted for the majority of intestinal IGLE in the duodenum (Fig.1G,H) confirming prior studies in rats (*5*). Conversely the RNG*^Cart^* neurons displayed few IGLE (Fig. 1G), preferentially innervating the villi and crypts of the duodenum (Fig1. F,I) as quantified by the significantly greater number of mucosal endings per mm than LNG*^Cart^* neurons (Fig1. J). These data illustrate the existence of a lateralized innervation and terminal morphology patterns of vagal sensory fibers in the intestine.

### Asymmetric sensing of meal-born signals

Based on transcriptomics and anatomical findings (Figure 1), we hypothesized that LNG and RNG neurons are responsive to different sensory modalities. To test this in live animals, we generated *Cart-GCaMP6s* mice in which the fluorescent, calcium indicator GCaMP6s is selectively expressed in *Cart-Cre* neurons allowing *in vivo* visualization of NG*^Cart^*neuronal activity in response to mechanical or chemosensory intestinal stimuli. Two photon microscopy was used for 4D volumetric imaging of NG neuron activity (x, y, z over time) in response to metabolic stimuli administered to the first 2cm of the intestine (Fig. 2A). In our preparation, the neurons were viable, exhibited baseline firing activity, and the same neurons were responsive to repeated peripheral stimuli (Fig. S3A-C and supplementary Video 1).

**Figure 2.**
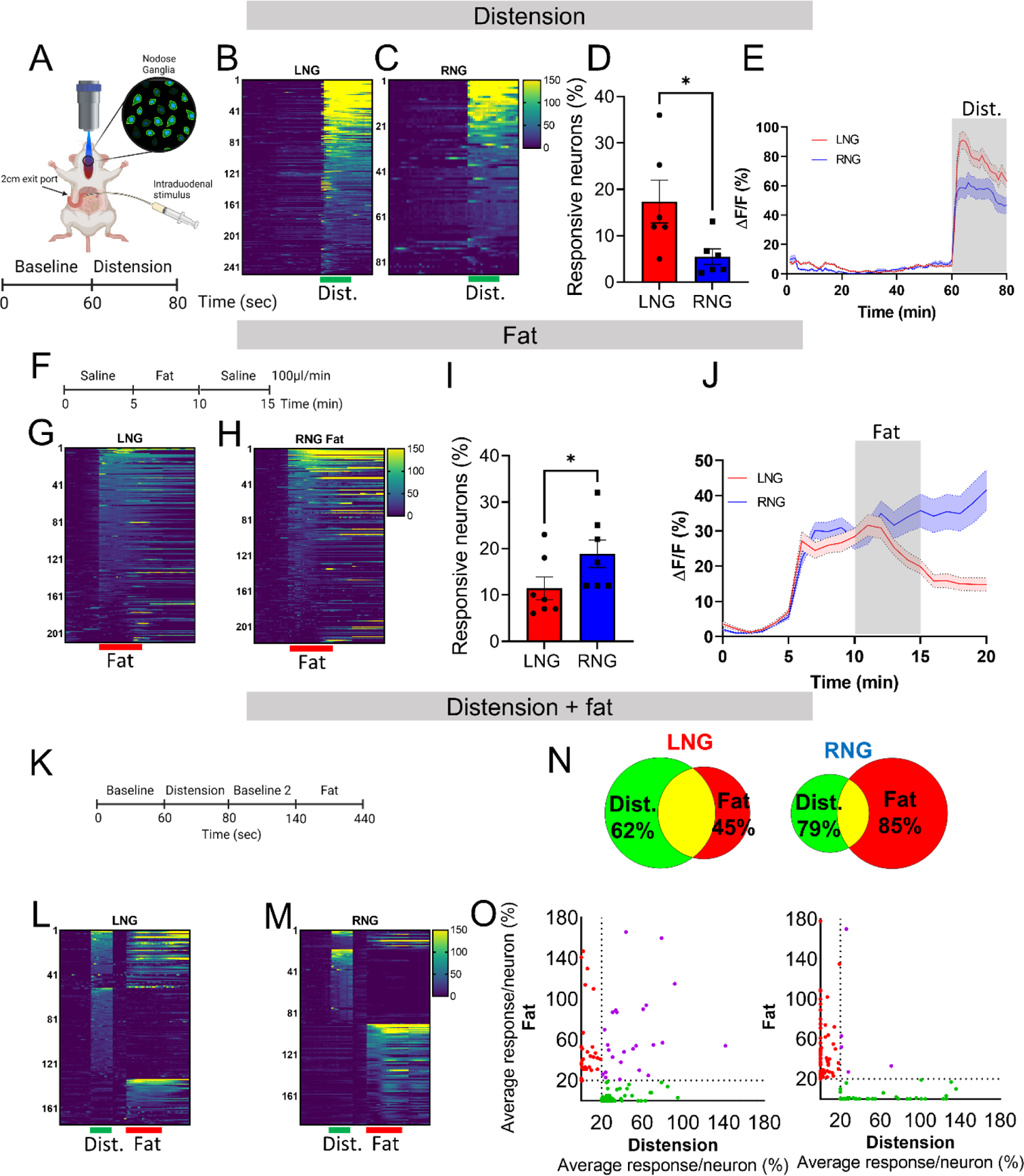
Activity of vagal sensory neurons in response to intraduodenal nutrient infusion or distension. (A) In vivo calcium imaging of NG*^Cart^* neurons in response to intestinal distension. (B-C) Heat maps depicting time-resolved responses (ΔF/F) of 250 LNG*^Cart^* and 85 RNG*^Cart^* neurons identified as distension responders (green bar, 20 seconds). (D) Percentage of NG*^Cart^* neurons identified as distension responders in LNG and RNG (n=6/group, unpaired t-test, *p=0.041). (E) Average GCaMP6s signal in LNG*^Cart^*and RNG*^Cart^* neurons that were responsive to duodenal distension. Grey shaded area represents duration of stimulus. Dark lines represent means and lighter shaded areas represent SEM. Two-way ANOVA, *p<0.0001. (F) In vivo calcium imaging of NG*^Cart^* neurons in response to intraduodenal fat infusion. (G-H) Heat maps depicting time-resolved responses (ΔF/F) of 210 LNG*^Cart^* and 219 RNG*^Cart^* neurons identified as fat responders (red bar, 5 minutes). (I) Percentage of NG*^Cart^*neurons identified as fat responders in LNG and RNG (n=7/group, Unpaired t-test, *p=0.049). (J) Average GCaMP6s signal in LNG*^Cart^* and RNG*^Cart^* neurons that were responsive to intraduodenal fat infusion. Grey shaded area represents duration of stimulus. Dark lines represent means and lighter shaded areas represent SEM. Two-way ANOVA, *p<0.0001. (K) In vivo calcium imaging of NG*^Cart^* neurons in response to intestinal distension and fat infusion. (L-M) Heat maps depicting time-resolved responses (ΔF/F) of 175 LNG*^Cart^* and 188 RNG*^Cart^* neurons identified as distension and/or fat responders (green bar, 20 seconds and red bar, 5 minutes). (N) Quantification of (L-M), n = 3 mice/group. (O) Average GCaMP6s signal in LNG*^Cart^*and RNG*^Cart^* neurons for the entire duration of the intraduodenal stimulus. Green dots= distension responsive; red dots = fat responsive; purple dots = responsive to both stimuli. Data are expressed as mean ± SEM; ns, p > 0.05; t tests and post hoc comparisons, *p < 0.05.

Air-induced duodenal distension (*19*), elicited rapid and robust calcium responses in NG*^Cart^* neurons (Fig. 2B,C). Distension preferentially recruited LNG*^Cart^* neurons (17.35%) compared to RNG*^Cart^*neurons (5.5%; Figures 2D, S3D) and led to a larger magnitude of response in LNG*^Cart^*neurons (Fig. 2E and S3E). NG*^Cart^*neuron activity was also increased in response to intestinal fat infusion, administered by means of a catheter surgically implanted through the pylorus with an exit port that allowed infusion over the first 2 cm of the intestine. Comparative analysis revealed that intraduodenal fat resulted in larger, more sustained activity in a greater percentage of RNG*^Cart^*(18.83%) compared to LNG*^Cart^* neurons (12.05%) (Fig. 2F-J and S3F,G). Consistent with previous reports of “salt-and-pepper” organization of NG (*7, 19*), the neurons were intermingled with no obvious topographical pattern.

The timescale and magnitude of vagal neuron response was different across sensory modalities, with faster and greater magnitudes occurring in response to mechanical, compared to nutritive, stimuli. Given the rapid reduction in neural activity in LNG neurons after termination of fat infusion compared to RNG (Fig. 2J and S3G), we hypothesized that LNG*^Cart^* neurons may be not responding to nutrient but rather to the fat-induced distension inherent with the infusion technique. To test this, we compared neuronal activity in response to both duodenal distension and fat infusion in the same animals (Fig. 2K-M). The majority of LNG*^Cart^*neurons that produced strong and continuous average activity in response to fat were also activated by intestinal stretch, while only a small fraction of RNG*^Cart^* neurons had sustained response to both stimuli (Fig. 2N-O and S3H-I). Thus, LNG*^Cart^*neurons are predominantly responsive to intestinal distension, while the RNG*^Cart^* population more abundantly sense intestinal fat. Altogether, these data provide functional evidence for the existence of a previously unsuspected asymmetry in vagal sensing of separate sensory modalities from intestinal-derived signals.

### Right vagal sensory neurons control food choice

Post-ingestive nutrient reinforcement can influence food preferences even without orosensory inputs (*20–22*), and this effect is mediated by vagal sensory neurons of the right NG (*23*). Here we tested whether a genetically-defined subpopulation of neurons that express CART are necessary and sufficient for flavor-nutrient conditioning. To test necessity, we implanted intragastric catheters in *Cart-Cre* mice and ablated RNG*^Cart^* neurons using flex-taCasp3 TEVp virus. Next, we performed a flavor-nutrient conditioning test in which novel non-caloric flavors were paired with either an intragastric delivery of fat (Microlipid, 7.5%) or saline (Fig. 3A). After 6 days of conditioning sessions, the control group significantly increased the preference for the fat-paired flavor, but fat reinforcement was impaired in animals with RNG*^Cart^*ablation (Fig. 3B). These results demonstrate that RNG*^Cart^* neurons are necessary for fat reinforcement. To test whether stimulation of RNG*^Cart^* neurons would be sufficient to promote flavor-conditioning, we targeted viral-mediated expression of the excitatory designer chemogenetic receptor hM3D(Gq) to either LNG*^Cart^* or RNG*^Cart^* neurons and performed a flavor conditioning test pairing non-nutritive flavors with either CNO or saline (Fig. 3C). Stimulation of RNG*^Cart^*, but not LNG*^Cart^*, neurons significantly increased preference for the CNO-paired flavor (Fig. 3D), demonstrating that RNG*^Cart^* neurons are sufficient for flavor reinforcement. To assess whether activation of these neurons is rewarding, we performed a self-stimulation experiment in mice with unilateral injection of the light-sensitive depolarizing channel *Channelrhodopsin2* (ChR2) (Fig. 3E). Mice with optogenetic posts implanted over vagal terminals in the NTS were placed in an operant chamber with an active and inactive nose hole in which nose pokes of the active hole would optically excite NTS terminals of LNG*^Cart^* or RNG*^Cart^* neurons (Fig. 3F). The mice expressing ChR2 in RNG*^Cart^* neurons learned to produce operant responses for optogenetic stimulation resulting in greater numbers of active nose pokes than LNG*^Cart^* mice that failed to engage in self-stimulation behavior (Fig. 3G). Additionally, we performed patch clamp recordings from dopamine neurons in substantia nigra pars compacta brain slices collected 30 minutes after intestinal infusion of saline, fat, or intestinal distension. Neural firing of dopamine neurons in the SNc was increased in response to fat compared to saline or distension treatment (Fig. S4). Altogether, these results indicate that RNG*^Cart^*neurons are necessary and sufficient for gut-brain reward that reinforces preference for foods with nutritive value.

**Figure 3.**
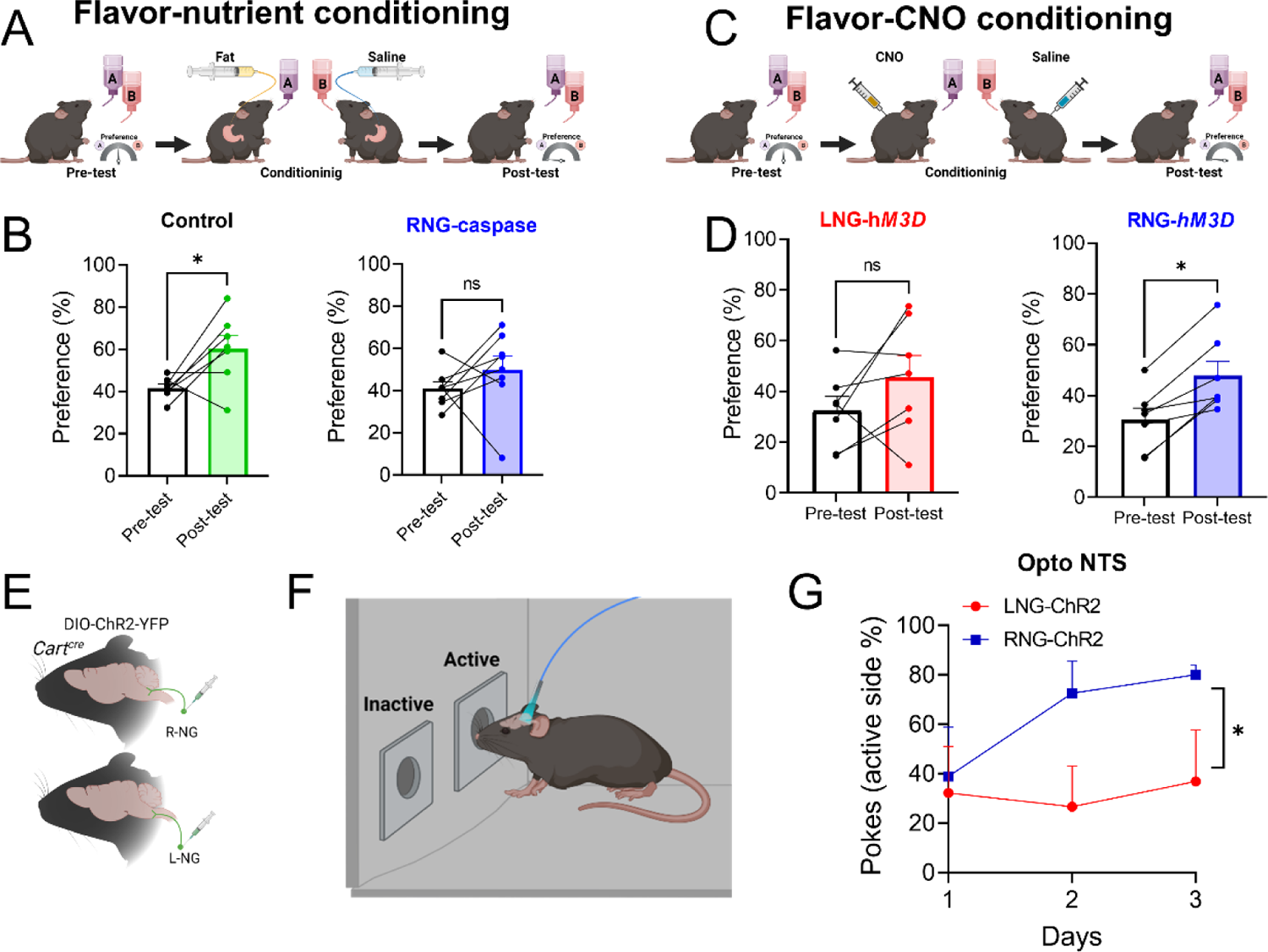
Stimulation of RNG*^Cart^* neurons promotes flavor-conditioning. (A) Flavor nutrient-conditioning test with mice that had RNG*^Cart^* neurons ablated with flex-taCasp3-TEVp. (B) Post-conditioning preferences for controls, and RNG*^Cart^*-Casp (n=6/group, paired t test). (C) Flavor CNO-conditioning test with LNG*^Cart^*-hM3Dq and RNG*^Cart^*-hM3Dq mice. (D) Post-conditioning preferences for LNG*^Cart^*-hM3Dq mice, and RNG*^Cart^*-hM3Dq mice (n=5-7/group, paired Student’s t test). (E) *Cart-Cre* mice unilaterally injected in the LNG or RNG with DIO-ChR2-YFP. (F) Mice implanted with optogenetic posts above vagal terminals in the NTS were trained to perform an operant task using a two-hole nose poke apparatus. One of the nose holes was designated as the active hole and was associated with a blue light laser pulse triggered by nose pokes, while the other hole remained inactive. (G) Preferences for the laser-paired nose poke over 3 consecutive days of stimulation for LNG*^Cart^*-ChR2 and RNG*^Cart^*-ChR2 mice (n=3-4/group, paired Student’s t test). Data are expressed as mean ± SEM; ns, p > 0.05; t tests and post hoc comparisons, *p < 0.05.

### Left vagal sensory neurons control stretch-induced satiety

Vagal gut-brain signaling plays a crucial role in the control of food intake through the coordinated actions of meal termination and satiety mechanisms (*24*). Importantly, genetically-defined subpopulations of vagal sensory neurons associated with mechanosensation can effectively elicit acute reductions in food intake, while stimulation of chemosensory populations has no impact on the amount of food consumed (*9*). To investigate the sufficiency of LNG*^Cart^* neurons in regulating food intake, we used unilateral chemogenetic stimulation to activate NG neurons in *Cart-Cre* mice injected with AAV9-hSyn-DIO-hM3D(Gq)-mCherry (Fig. 4A-D). We observed a 30-40% reduction in food intake that persisted for 6 hours following stimulations of LNG*^Cart^*neurons, with the greatest effects observed during the first 4 hours after CNO injection (Fig. 4E-F, and S5A,B). This reduction was exclusively due to decreased meal size, as there was no effect on meal duration or number (Fig. 4F and S5E). Notably chemogenetic stimulation of RNG*^Cart^*neurons had no effect on acute food intake (Fig. 4 G,H and S5C-F). To gain insight into the mechanism underling LNG*^Cart^* neuron induced satiation we tested gastric emptying in response to chemogenetic stimulation using an acetaminophen assay (Fig. 3I). We found that stimulation of LNG*^Cart^* neurons led to reduced circulating acetaminophen levels compared to stimulation of RNG*^Cart^* neurons or in control mice (Fig. 4J), suggesting that recruitment of LNG*^Cart^* neurons delays gastric emptying.

**Figure 4.**
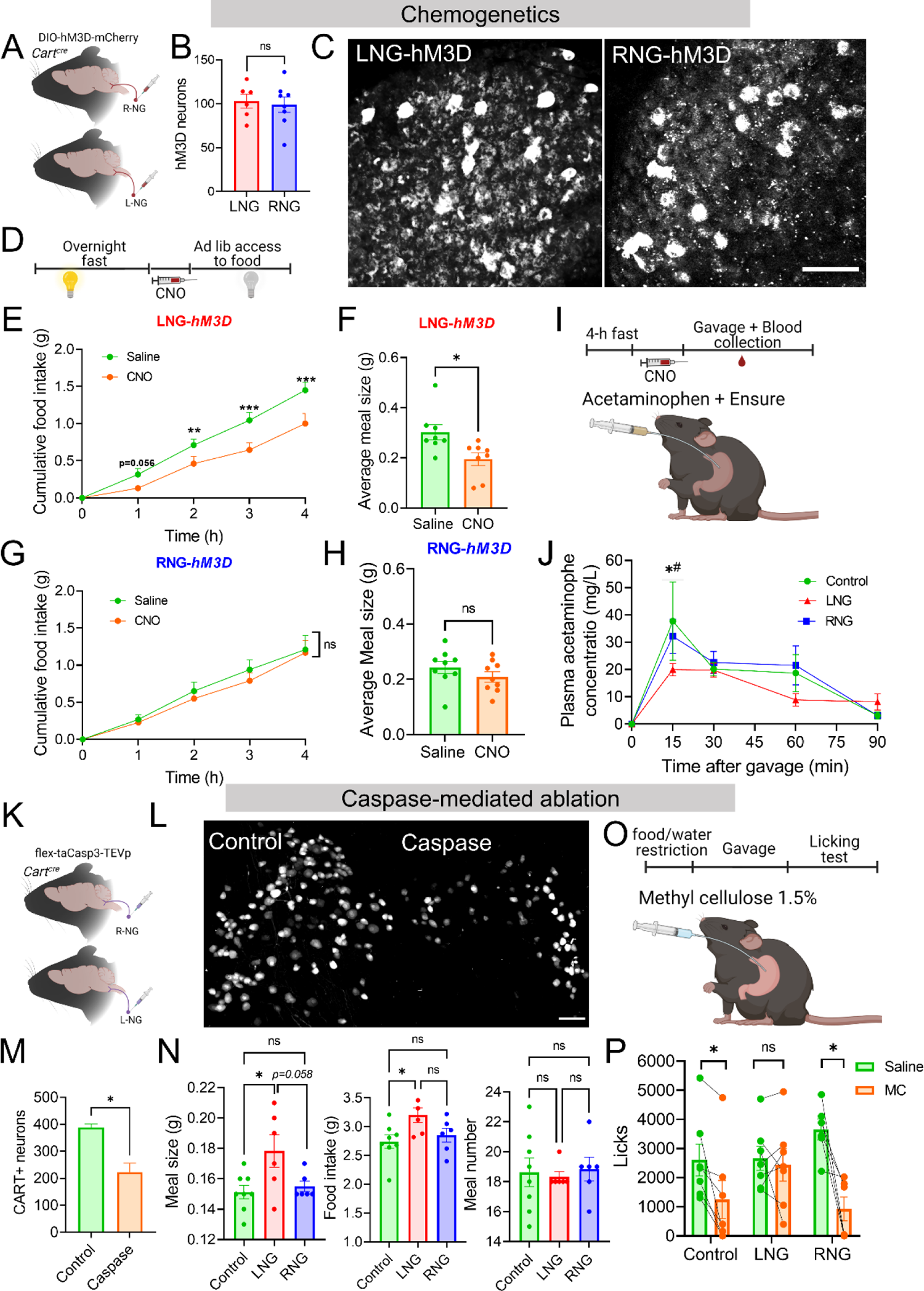
Stimulation of LNG*^Cart^* neurons is sufficient and necessary to reduces food intake. (**A**) *Cart-Cre* mice unilaterally injected in the LNG or RNG with DIO-hM3D(Gq)-mCherry. (**B**) Quantification of hM3D-mCherry expression in LNG and RNG. (**C**) hM3D-mCherry detected in LNG and RNG. (**D**) Paradigm for temporal analysis of chow intake in LNG*^Cart^*- and RNG*^Cart^*-hM3Dq mice. (**E-H**) 4-h food intake of (E) LNG*^Cart^*-hM3Dq mice, and (G) RNG*^Cart^*-hM3Dq mice (n=9/group, two-way ANOVA with Two-stage linear step-up procedure of Benjamini, Krieger and Yekutieli post hoc) following CNO or saline injections; Average 4-h meal size of (F) LNG*^Cart^*-hM3Dq, and (H) RNG*^Cart^*-hM3Dq mice (n = 9/group, paired t test, ns) following CNO or saline injections. (**I**) Blood collection for gastric emptying assay from LNG*^Cart^*- and RNG*^Cart^*-hM3Dq mice following CNO injection and intragastric delivery of acetaminophen solution. (**J**) Gastric output measured as blood acetaminophen levels over time in LNG*^Cart^*- and RNG*^Cart^*-hM3Dq mice (n=4-8/group, two-way ANOVA with Two-stage linear step-up procedure of Benjamini, Krieger and Yekutieli post hoc; Control vs. LNG, *p = 0.018; RNG vs. LNG, #p = 0.033). (**K**) *Cart-tdTomato* mice unilaterally injected in the LNG or RNG with flex-taCasp3-TEVp. (**L**) NG from *Cart-tdTomato* mice after caspase-mediated ablation. (**M**) Quantification of (L). (**N**) 3-day average meal pattern parameters of ad lib, chow-fed LNG*^Cart^*- and RNG*^Cart^*-Casp mice (n = 6-8/group, one-way ANOVA with Holm-Sidak post hoc). (**O**) Intragastric delivery of methyl cellulose or saline to food and water restricted LNG*^Cart^*- and RNG*^Cart^*-Casp mice before licking test. (**P**) Intake of intralipid from LNG*^Cart^*- and RNG*^Cart^*-Casp mice following gavage of saline or methyl cellulose, (n = 5-6/group, paired Student’s t test). Data are expressed as mean ± SEM; ns, p > 0.05, *p < 0.05, **p < 0.01, ***p < 0.001. Scale bars 100 µm.

To assess the necessity of LNG*^Cart^*neurons for regulating food intake, we selectively ablated these neurons using a viral-mediated caspase in Cart-Cre mice (Fig. 4K-M). We found that elimination of LNG*^Cart^* neurons led to an increase in food intake that was driven by increased meal size, while ablation of RNG*^Cart^* neurons had no effect on food intake (Fig. 4N and S5G). To further investigate the role of LNG*^Cart^* neurons in meal termination, we assessed the effect of distension on food intake in mice with ablated LNG*^Cart^*neurons. We found that preloading mice with methylcellulose, a distension-inducing agent, led to reduction in fat intake in control and RNG*^Cart^*neurons, but not in mice with ablated LNG*^Cart^* neurons (Fig. 4O,P). Collectively, these findings demonstrate that LNG*^Cart^* neurons, but not RNG*^Cart^*neurons, are necessary and sufficient for satiation.

### Functional specialization of gut-brain circuits

Next, we aimed to investigate how different sensory inputs from the gut are integrated by the brain. To investigate whether gut-derived stimuli result in distinguishable patterns of neuronal activity we used FosTRAP mice to compare neural activity (*20, 25*) by analyzing tdTomato labeled TRAP neurons in response to intragastric fat infusion and cFos positive neurons in response to duodenal distension at two different timepoints in the same mouse (Fig. 5A). In the dorsal vagal complex (DVC), the first site of vagal sensory integration (*17*), we found that both intragastric fat infusions and intestinal distension resulted in extensive neuronal labeling (Fig. 5B-D), with similar number of total Fos positive neurons (Fig. 5B). However, there was significant separation in neuronal activity between fat infusion and intestinal distension within the DVC neurons between groups (Fig. 5C-D). Specifically, we find that only 26% of the DVC neurons are responsive to both fat infusions and distension (Fig. 5C). Interestingly, fat resulted in a greater proportion of neuronal labeling in the NTS, while distension resulted in a greater fraction of active DMV neurons (Fig. 5B). Interestingly, selective optogenetic stimulation of duodenal terminals from unilateral NG neurons results in similar shifts in NTS/DMV activity (Fig. 5E-H). ChR2 labeled vagal terminals came into close apposition with cFos labeled post-synaptic NTS neurons (Fig. S6A). The preferential recruitment of distension-sensing LNG*^Cart^* neurons of a vago-vagal reflex is consistent with evidence that LNG*^Cart^*neurons slow gastric emptying. Altogether, these results highlight functionally specialized hindbrain circuits in response to mechanical and chemical stimuli is conveyed by asymmetrical gut-brain mechanisms.

**Figure 5.**
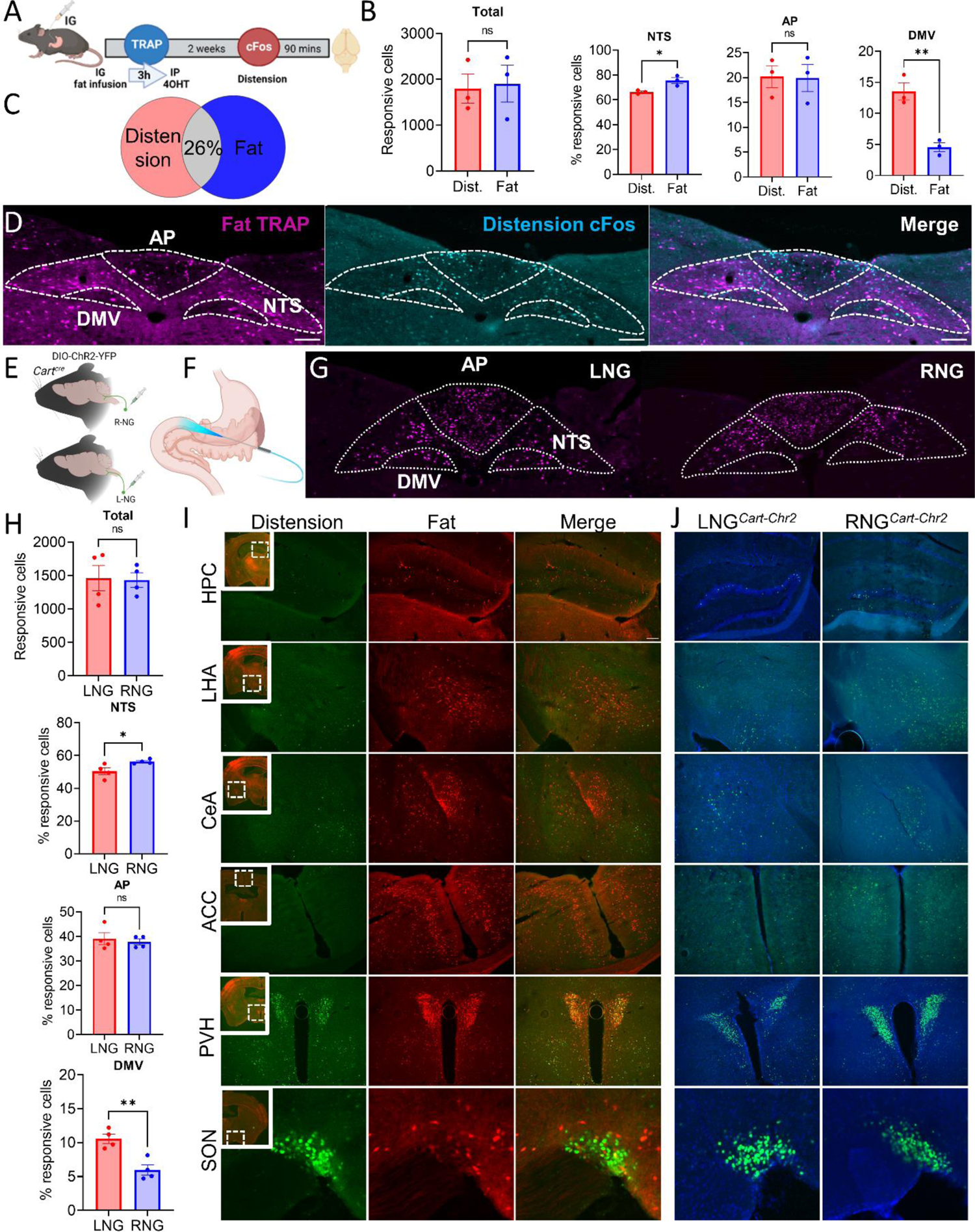
LNG*^Cart^* and RNG*^Cart^* recruit distinct downstream circuits in the brain. (**A**) Schematic of of the FosTRAP model for comparing neural response to intraduodenal fat and distension in the same animal. (**B**) Quantification of tdTomato labeled FatTRAP neurons and neurons labeled with Fos immunoreactivity after duodenal distension in the DVC (unpaired Student’s t test) (**C**) Venn diagram representing the separation in neuronal activity between the two stimuli. (**D**) Representative images of the NTS showing active populations in response to fat (magenta) and distension (cyan). (**E**) Unilateral injection of DIO-ChR2-YFP. (**F**) Optogenetic stimulation of the duodenum. (**G**) Representative images of the NTS after optogenetic stimulation of LNG*^Cart^* or RNG*^Cart^* fibers that innervate the duodenum. (**H**) Quantification of cFos expression on the DVC 90 min after stimulation (unpaired t test). (**I**) Brain-wide screen of FosTRAP mice that received intestinal distension and fat infusion. (**J**) Brain-wide screen of LNG*^Cart^*-ChR2 or RNG*^Cart^*-ChR2 mice that received intraduodenal optogenetic stimulation. HPC=hippocampus; LHA=lateral hypothalamus; CeA=Central amygdala; ACC=anterior cincular cortex; PVH=paraventricular nucleus of the hypothalamus; SON=supraoptic nucleus. Data are expressed as mean ± SEM; ns, p > 0.05; t tests and post hoc comparisons, *p < 0.05. Scale bars 100 µm.

In order to start defining central circuits that are recruited by different interoceptive stimuli, we performed a brain-wide screen of FosTRAP mice following intestinal distension or fat infusion and a detailed description of expression patterns is provided in supplementary fig. 6D. We observed crosstalk between calories and food volume in the paraventricular nucleus (PVH, Fig. 5I), and modest levels of overlap in the paraventricular nucleus of the thalamus (PVT, Fig. S6B). We confirmed that these brain regions are activated in response to optogenetic stimulation of left and right NG*^Cart^* neurons innervating the intestine (Fig. 5J and S6C). It was also notable that the ventromedial hypothalamus (VMH) had no labeling in response to either stimuli or optogenetic stimulation of duodenal-innervating NG*^Cart^* neurons, suggesting that the VMH is not responsive to vagal interoceptive signals irrespective of sensory modality (Fig. S6B,C). Of particular interest, distinct activation patterns were observed in response to distension vs fat. Distension engaged brain regions associated with energy homeostasis, including the arcuate nucleus, dorsal medial hypothalamus, supraoptic nucleus, and ventral bed nucleus of the stria terminalis (vBNST). Conversely, fat infusion predominantly activated brain regions associated with salience and cognition, such as the anterior cingulate cortex, hippocampus, lateral hypothalamus, dorsal striatum, amygdala, and dorsal BNST (Fig. 5I, and S6B). Optogenetic stimulation of vagal terminals in the duodenum supported a role for RNG*^Cart^* neurons in nutrient activated brain regions associated with cognition, while LNG*^Cart^* neurons signal information about distension to brain regions associated with energy balance (Fig. 5J, and S6C). In summary, meal-related information from the gut is broadly integrated by many brain regions, but discrete nuclei were identified that detect singular interoceptive modalities via asymmetric gut-brain circuits.

## DISCUSSION

Altogether, these findings unravel the different roles of left and right gut-brain circuits in various aspects of feeding. Our research reveals that a subset of genetically-defined gut-innervating vagal sensory neurons expressing the neuropeptide CART are highly lateralized in respect to their gene expression, anatomy, sensory response to intestinal stimuli, behavioral consequence, and recruitment of central circuits. This work provides new insight into how interoceptive information about the volume and quality of food is resolved by distinct lateralized gut-brain circuits to enable rapid eating decisions.

Distension of the gastrointestinal tract conveys information about food volume, and serves as a key mechanism that controls food intake. A critical role for gastric distension in reducing food intake is supported by studies that use gastric balloons to inflate the stomach (*26*), or pyloric cuffs that retain food within the stomach (*27*). Vagal sensory neurons with well-defined morphological terminals (*28*) and genetic markers (*9, 19, 29*) are activated by inflation of gastric balloons (*30*) and these neuronal populations recruit central circuits (*31–33*) capable of reducing food intake (*9, 31, 33*). Loss of function experiments using subdiaphragmatic or selective branch vagotomies (*34, 35*) suppress reductions in food intake in response to gastric distension, suggesting that the vagus nerve is necessary for gastric distension-induced satiety. Yet studies targeting gastric vagal sensory neurons more precisely (*36, 37*) have questioned their necessity in appetite suppression.

Recently, separate vagal sensory neurons were identified to response to gastric or intestinal distension (*9, 19, 29*), and stimulation of intestinal vagal mechanosensory populations more potently inhibited food intake than vagal populations that primarily innervate the stomach (*9*). Our findings demonstrate the existence of a distinctive subpopulation of CART-expressing neurons in the LNG that possess exquisite sensitivity to intestinal stretch and prominently innervate the duodenal muscular layer. Stimulation of these LNG*^Cart^* neurons slows gastric emptying and inhibits food intake. Although stimulation of intestinal terminals of these neurons recruits a vago-vagal reflex that can influence gastric emptying and suppression of food intake (*38*), LNG*^Cart^* neurons also innervate the stomach (*16*) so further research is required to tease apart the individual contributions of gastric and intestinal mechanosensation in modulating satiety.

We previously identified a circuit that connects the gut to nigrostriatal dopamine neurons and signals gut reward via the RNG (*23*). Here we identify CART as a genetic marker that defines the subset of RNG neurons responsible for self-stimulation, a hallmark behavior of reward. RNG*^Cart^* neurons co-express molecular markers associated with chemosensation and extensively innervate villi and crypts, making them ideally positioned to sense nutrients that are absorbed from the intestinal lumen. We demonstrate that these neurons are activated in response to intestinal fats and are capable of increasing food preferences over multiple trials, however they do not result in acute changes in the amount of food consumed. These data suggest that asymmetrical vagal circuits engage different sets of behaviors that influence food intake over different timescales.

Optimizing food choice is based on the ability to learn, remember, and predict the rewarding value of a food based on prior experience. In addition to the recruitment of nigrostriatal circuits, we identify four novel chemosensory gut-brain circuits mediated primarily by the right NG involved in cognition. The hippocampus, a region associated with learning and memory (*39*), receives interoceptive information from gut-innervating vagal sensory neurons (*40*). Here we expand these findings by demonstrating asymmetrical recruitment of a gut-hippocampal circuit in response to nutritive stimuli sensed by chemosensory RNG neurons. Secondly, we demonstrate that stimulation of intestinal RNG terminals and infusions of fat selectively activate the central amygdala, a brain region that influences learning and motivated feeding behavior related to sensory information (*41–43*). Furthermore, the lateral hypothalamus, a brain region necessary for reinforcement learning (*44*) that links motivation with cognitive processes (*45*), was preferentially recruited by fat and RNG stimulation. Finally, we identify the anterior cingulate cortex as another area that preferentially integrates nutritive, over mechanosensory, information from the gut. The anterior cingulate cortex has been associated with preparatory, and/or maintenance of attention that would facilitate cognition in response to interoceptive signals (*46, 47*), thereby creating a psychological state that would enhance learning in response to gut reward. Disruption of RNG*^Cart^*signaling could reduce flexibility to adapt to post-ingestive cues, and instead lead to habitual eating patterns. Thus, RNG*^Cart^* neurons could be a target for improving cognitive flexibility.

Lateralization of neural circuits is evolutionarily conserved. Fish use lateralization to improve their ability to swim in unison (*48*). Birds use lateralized visual fields to improve feeding while monitoring for predators (*49*). C.elegans use left or right ASE neurons to sense food and engage motility (*11*). Human language processing is highly lateralized (*12*). It has been proposed that lateralization of neural processes evolved to increase integration of multiple sensory information (*12*). Defined genetic signatures underlying the specialized sensory responses of NG neurons have been previously described (*7*). Our findings identify asymmetrical gene expression in NG^Cart^ neurons, but also that this is a ubiquitous feature of NG neurons. Together with existing anatomical evidence for lateralized innervation patterns of the heart (*50*), liver (*51*), lungs (*52*), pancreas (*53*), and stomach (*5*) this suggests a common mechanism of sensory processing that encodes functional organization of vagal interoception. The implication of these findings for our understanding of the neural mechanisms that underlie interoceptive perception and their potential role in health and disease warrant further exploration. In light of recent advances in spatial targeting of the vagus nerve for bioelectric medicine (*54–57*) and non-invasive optogenetic stimulation of central (*58*) and peripheral targets (*59*), the current findings provide an opportunity for improved therapeutic effectiveness of vagal nerve stimulation for treating metabolic and neuropsychiatric diseases.

## Author contributions

### Data and materials availability

All materials are available upon request or are commercially available.

---

**Supplementary Figure 1.**
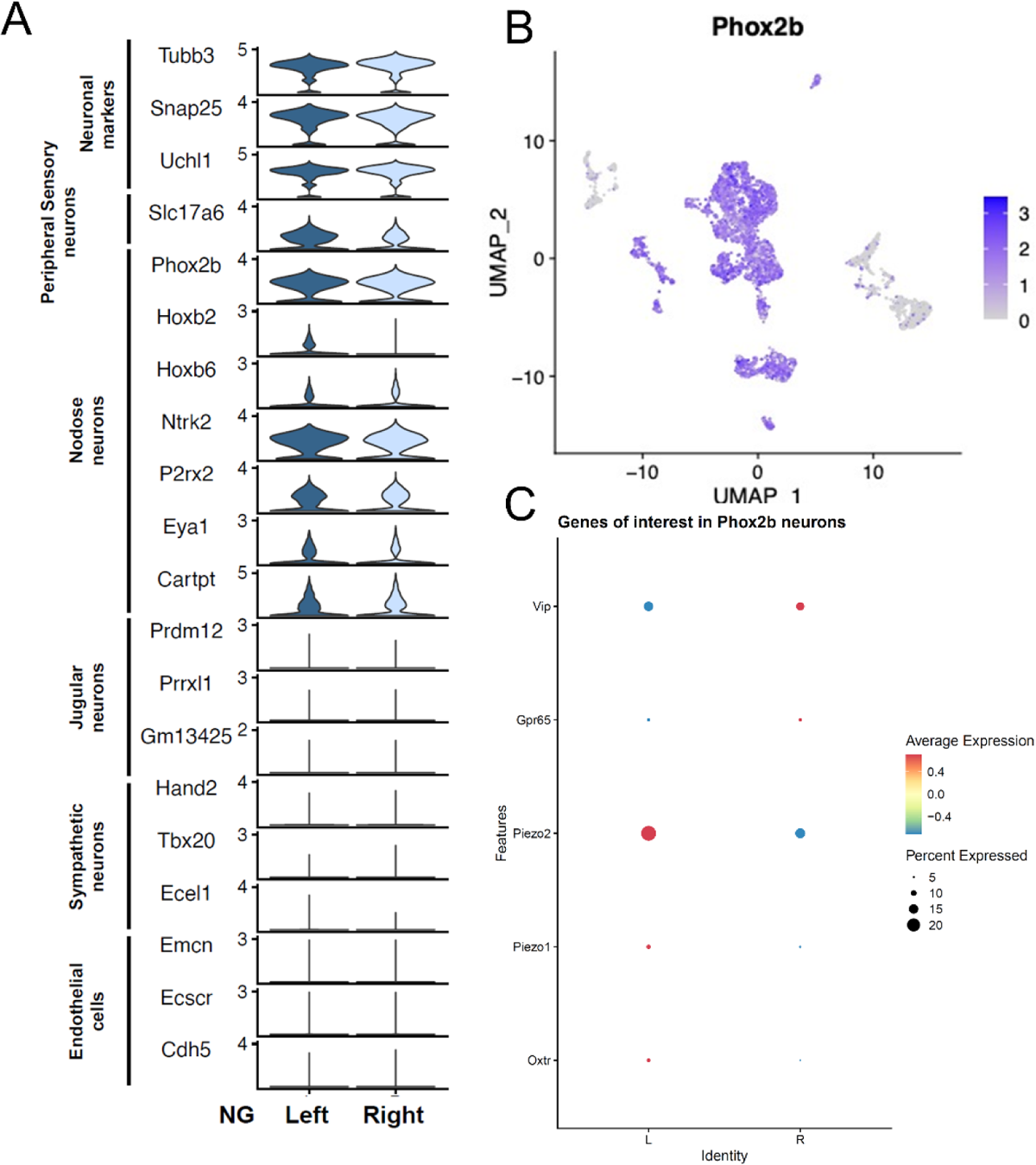
Single cell RNA sequencing data of LNG*^Cart^* and RNG*^Cart^*, related to figure 1. (**A**) Violin plots showing the expression of genes exclusively present in NG neurons. (**B**) UMAP illustrating the selectivity for *Phox2b* nodose placode-derived neurons in our analysis. (**C**) Molecular markers underlying the anatomical differences between LNG*^Cart^* and RNG*^Cart^*. The dot plot shows expression of selected genes involved in mechanosensing and nutrient sensing in all NG neurons.

**Supplementary Figure 2.**
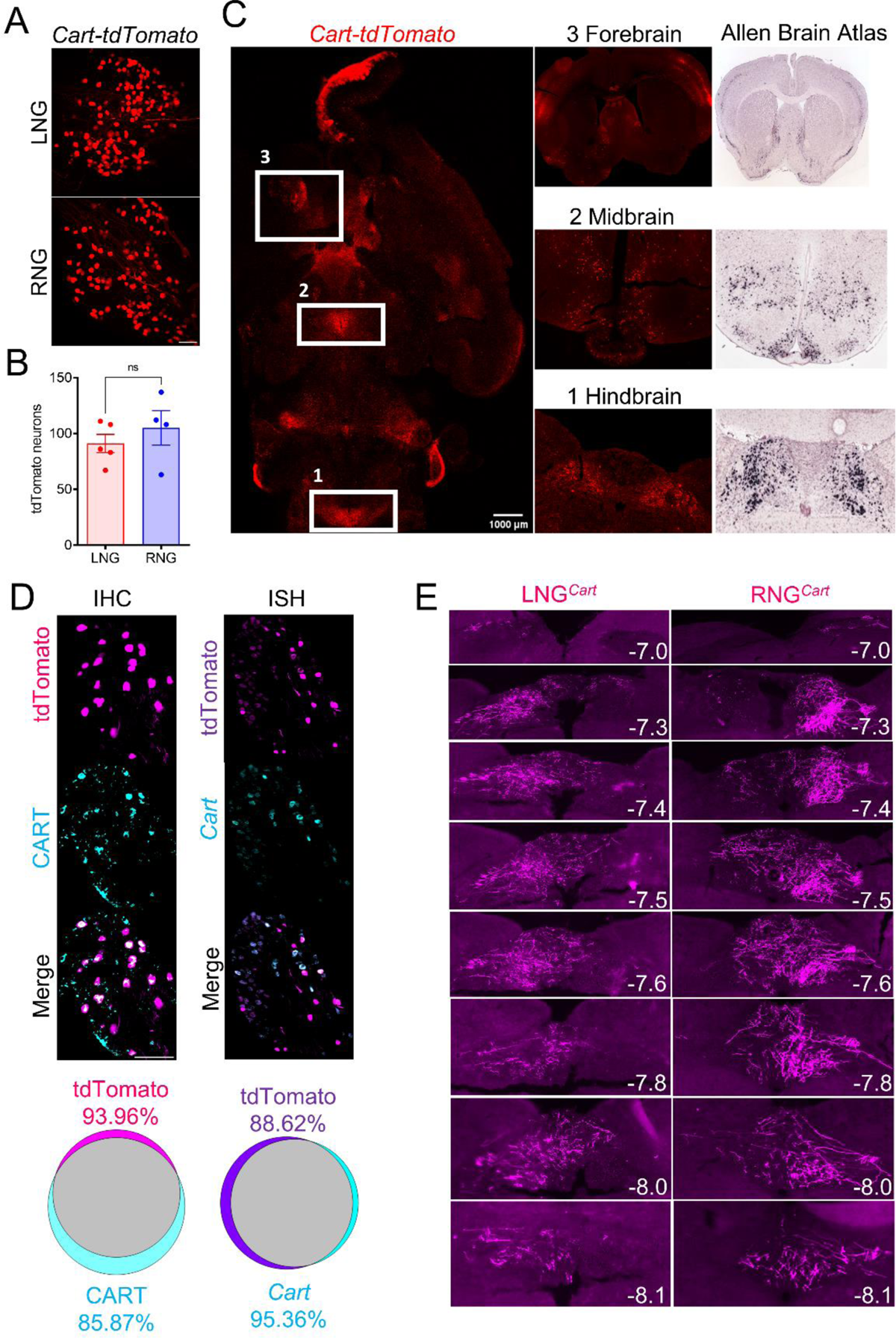
Validation of *Cart-Cre* mouse line, related to figure 1. (**A**) tdTomato detected in NG of *Cart-tdTomato* mouse line. (**B**) Quantification revealed similar number of CART neurons in both LNG and RNG. (**C**) Left: Axial brain section illustrating tdTomato labeling in *Cart*-expressing regions. Right: Coronal sections displaying tdTomato-positive neurons in brain areas known for *Cart* expression. Allen Brain Atlas was utilized for comparison purposes. (**D**) Immunohistochemistry and In Situ hybridization revealing the amount of overlap between Cart protein and mRNA and tdTomato labeling in the NG. (**E**) Brainstem sections showing the tdTomato+ axonal terminals of LNG*^Cart^* and RNG*^Cart^* in the dorsal vagal complex. AAV9-DIO-tdTomato was injected unilaterally into the nodose ganglion of *Cart-Cre* mice. Scale bars 100 µm.

**Supplementary Figure 3.**
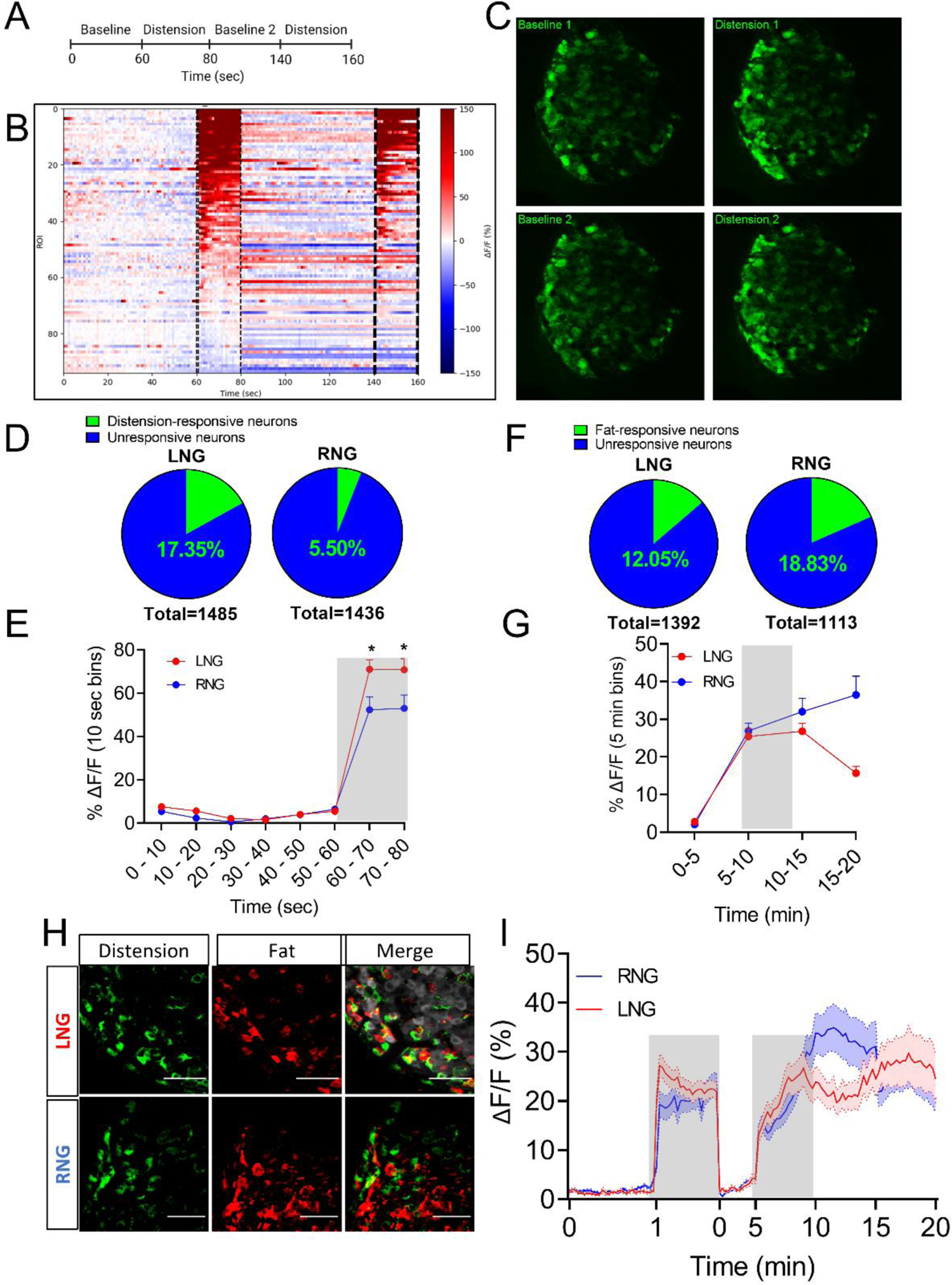
Activity of vagal sensory neurons in response to intraduodenal nutrient infusion or distension, related to figure 2. (A) In vivo calcium imaging of NG*^Cart^* neurons in response to repeated intestinal distension. (B) Heat maps depicting time-resolved responses (ΔF/F) of neurons in supplementary video 1. (**C**) Snapshots of supplementary video 1 showing that same population of NG*^Cart^* neurons responds to repeated distension stimulus applied to the duodenum. (**D**) Percentage of distension-responsive NG*^Cart^* neurons in LNG and RNG (n=6). (**E**) Average GCaMP6s signals (10 sec bins) in LNG*^Cart^* and RNG*^Cart^*neurons following duodenal distension. Grey shaded area represents duration of stimulus (n=6/group, two-way ANOVA, p=0.0004). (**F**) Percentage of fat-responsive CART neurons in LNG and RNG (n=7). (**G**) Average GCaMP6s signals (5 min bins) in LNG*^Cart^* and RNG*^Cart^* neurons following intraduodenal fat infusion. Grey shaded area represents duration of stimulus (n=6/group, two-way ANOVA, p<0.0001). (**H**) Images of Left and Right NG of *Cart-GCaMP6* mice showing calcium fluorescence responses to duodenal distension (green) and fat infusion (red). Scale bar, 100 µm. (**I**) Average GCaMP6s signals (10 sec bins) in LNG*^Cart^* and RNG*^Cart^* neurons in response to duodenal distension and fat infusion. Grey shaded area represents duration of stimuli (n=3/group). Data are expressed as mean ± SEM; ns, p > 0.05; t tests and post hoc comparisons, *p < 0.05.

**Supplementary Figure 4.**
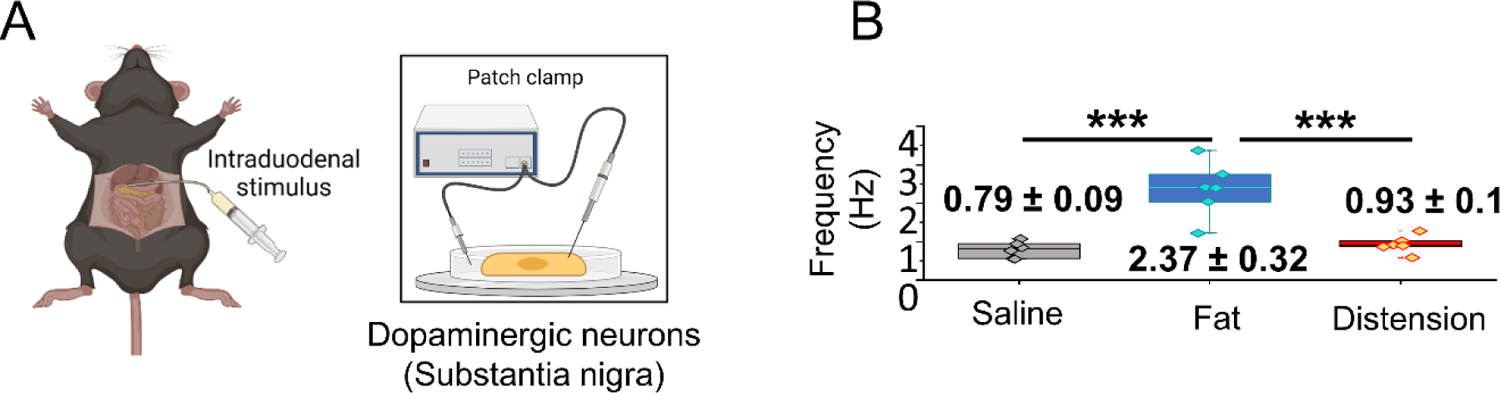
Stimulation of RNG*^Cart^* neurons promotes neural firing of dopamine neurons in the substantia nigra, related to figure 3. (**A-B**) Mice received intraduodenal stimulus and patch clamp electrophysiology was used to measure the firing rate of dopaminergic neurons of the substantia nigra. (I) firing rate of dopaminergic neurons of the substantia nigra (one-way ANOVA with Holm-Sidak post hoc). N denotes the number of neurons for in vitro experiments. Data are expressed as mean ± SEM; ns, p > 0.05, *p < 0.05, **p < 0.01, ***p < 0.001.

**Supplementary Figure 5.**
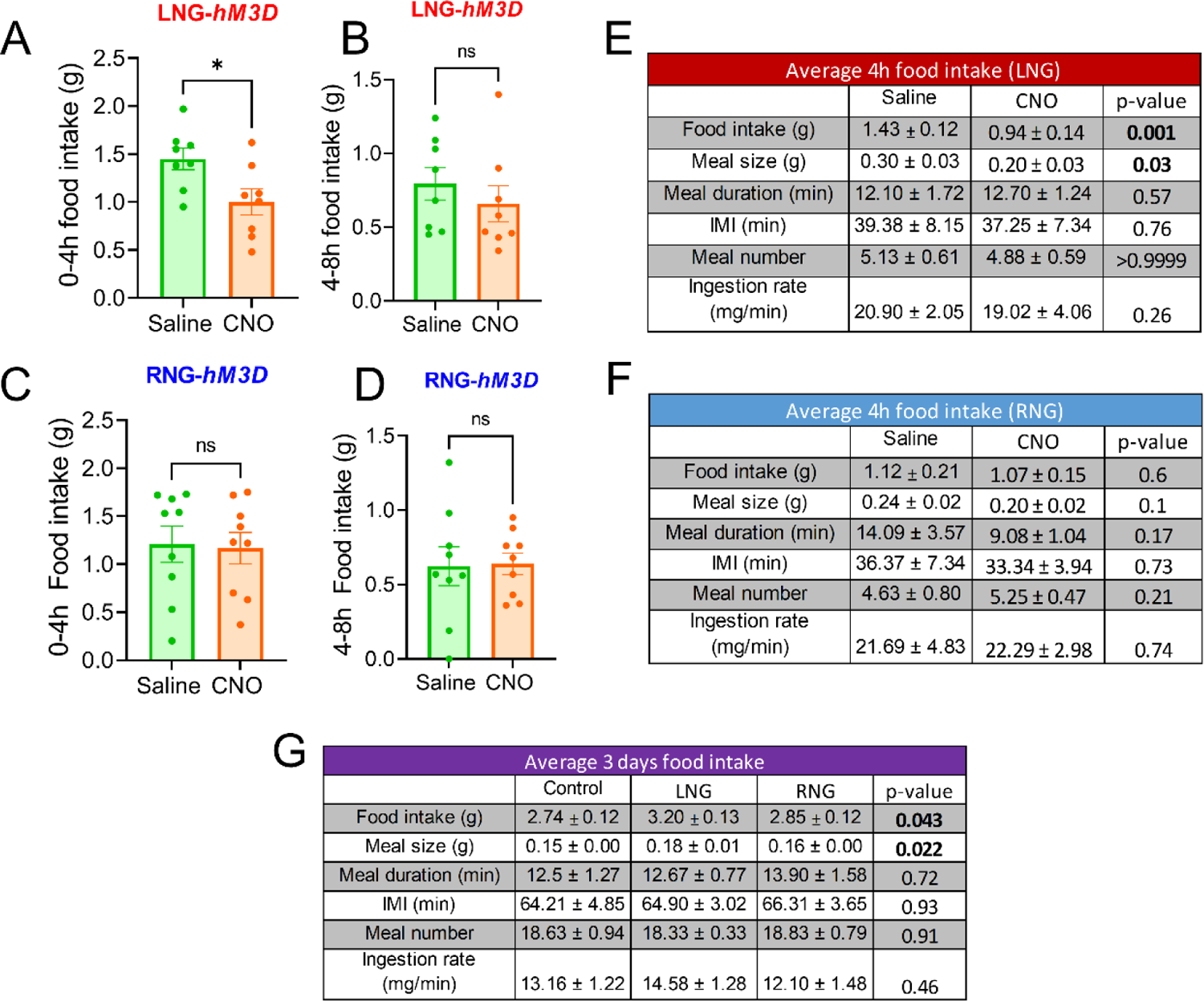
LNG*^Cart^* neurons regulate meal size, related to figure 4. (**A-D**) Cumulative food intake was analyzed in 4-h bins; Initial 4-h food intake of (A) LNGCart-hM3Dq, and (C) RNGCart-hM3Dq mice (n = 8/group, paired Student’s t test) following CNO or saline injections; Second 4-h bin of cumulative food intake; 4-8-h food intake of (B) LNGCart-hM3Dq, and (D) RNGCart-hM3Dq mice (n=8-9, paired Student’s t test) following CNO or saline injections. (**E-F**) Meal patterns of LNG*^Cart^*- and RNG*^Cart^*-hM3Dq mice were recorded averaging 4-h of food intake following CNO or saline injections in mice acclimated to BioDAQ (n = 9/group, paired Student’s t test). (**G**) Meal patterns of LNG*^Cart^*- and RNG*^Cart^*-Casp mice were recorded averaging three consecutive days of food intake in mice acclimated to BioDAQ (n = 6/group, one-way ANOVA with Holm-Sidak post hoc). Data are expressed as mean ± SEM; ns, p > 0.05, *p < 0.05, **p < 0.01, ***p < 0.001.

**Supplementary Figure 6.**
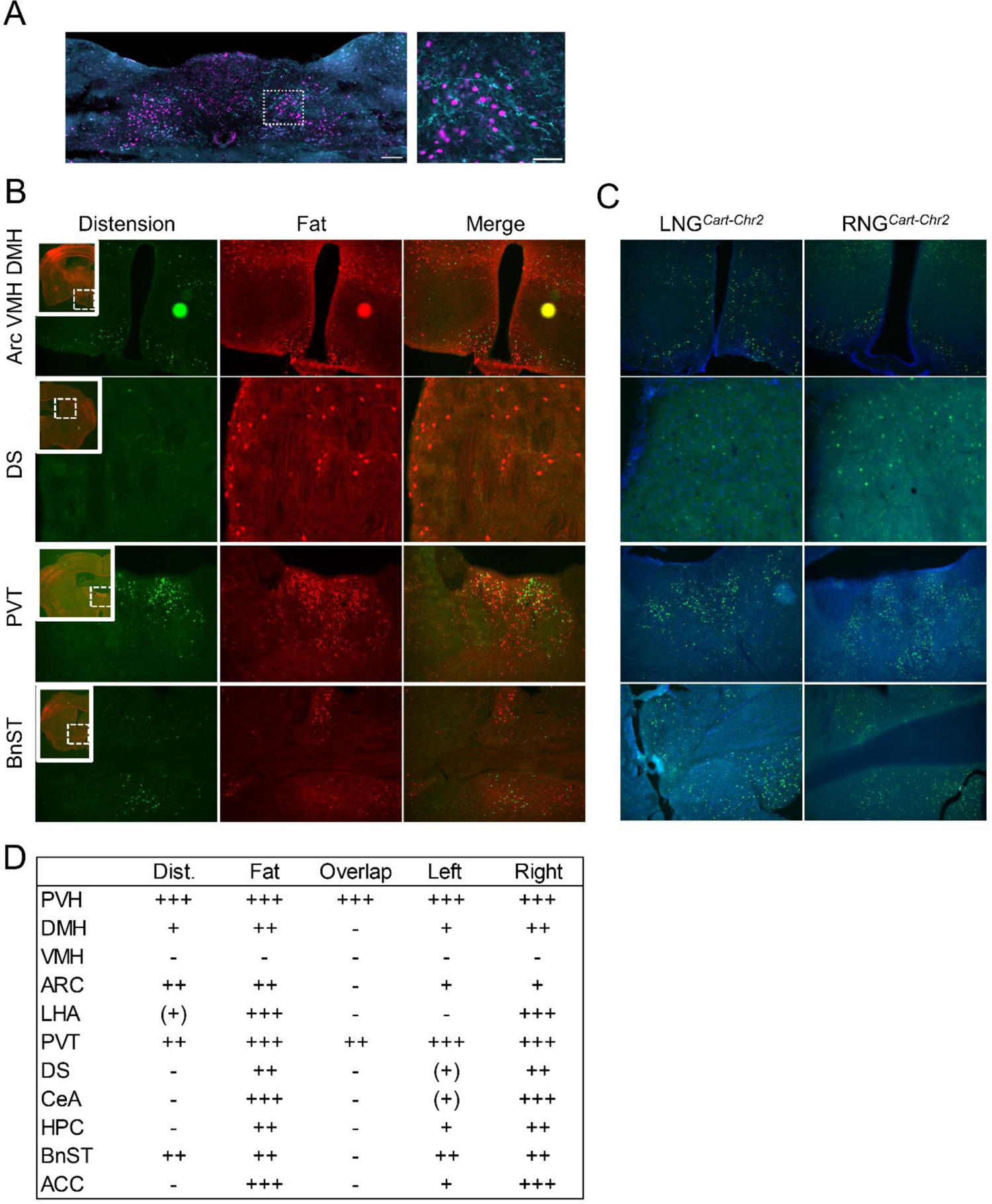
Distinct Activation Patterns of Central Circuits in Response to Intestinal Distension and Fat Infusion in FosTRAP Mice, related to figure 5. (**A**) Representative image of ChR2+ LNG*^Cart^* and RNG*^Cart^* vagal terminals in close proximity with cFos labeled post-synaptic NTS neurons. (**B**) Brain-wide screen of FosTRAP mice that received intestinal distension and fat infusion. (**C**) Brain-wide screen of LNG*^Cart^*-ChR2 or RNG*^Cart^*-ChR2 mice that received intraduodenal optogenetic stimulation. (**D**) Table with detailed description of cFos expression patterns in the brain, related to figures 5 I and J, and Sup. Fig. S6 B and C. Low, moderate and high expression patterns are shown as +, ++ and +++, respectively. Undetected and unclear expressions are shown as – and (+), respectively. Arc=arcuate nucleus VMH=ventral medial hypothalamus DMH=dorsal medial hypothalamus DS=dorsal striatum PVT=paraventricular nucleus of the thalamus BnST=bed nunclei of stria terminalis. Scale bars 100 µm.

## Materials and Methods

### Materials Availability

This study did not generate new unique reagents.

### Data and Code Availability

The published article includes all datasets generated or analyzed during this study. Detailed datasets and codes supporting the current study are available from the lead contact upon request.

### Animals

All mice used in experiments were individually housed under a 12 hr light/dark cycle in a room set to 23 °C (73 °F) with ad libitum access to food and water unless stated otherwise. Both male and female adult mice (at least 8 weeks old) were used, and we did not observe any sex differences. Mice were randomly assigned to experimental conditions and in a counter-balanced fashion with regards to sex and age. All procedures and experiments presented in this study were conducted in accordance to NIH guidelines and were approved by Animal Care and Use Committees from University of Florida and Monell Chemical Senses Center. Mouse strains details are as follows: C57BL/6J (Jax 000664), B6;129S-Cartpttm1.1(cre)Hze/J (Jax 028533), B6.Cg-Gt(ROSA)26Sortm14(CAG-tdTomato)Hze/J (Jax 007914), B6;129S-Gt(ROSA)26Sortm95.1(CAG-GCaMP6f)Hze/J (Jax 024105) B6.129(Cg)-Fostm1.1(cre/ERT2)Luo/J (Jax 021882), and Ai14 (B6.Cg-Gt(ROSA)26Sortm14(CAG-tdTomato)Hze/J, (Jax.007914).

### Viral vectors

All viral vectors were obtained from Addgene. PHP.S-FLEX-tdTomato (28306), AAV9-EF1a-double floxed-hChR2(H134R)-EYFP-WPRE-HGHpA (20298), AAV9-hSyn-DIO-hM3D(Gq)-mCherry (44361) and AAV5-flex-taCasp3-TEVp (45580).

### Surgical Procedures

#### Nodose Ganglia Injections

Mice received a subcutaneous injection of carprofen (5 mg/kg; Henry Schein) and underwent anesthesia (isoflurane,1.5-2.5%). Next, mice were placed on a heating pad on supine position and a 2 cm midline incision was made in the skin on the ventral aspect of the neck. The skin, salivary glands and underlying muscles were retracted, and the vagus nerve was blunt dissected from the carotid artery using fine-tip forceps. The NG was located by tracing the vagus nerve toward the base of the skull, and then exposed by retracting surrounding muscles and blunt dissection of connective tissues. A glass micropipette filled with viral construct attached to a micromanipulator was used to position and puncture the NG. A total of 0.5 µl was injected unilaterally into the NG using a Picospritzer III injector (Parker Hannifin, Pine Brook, NJ). Incision was closed and post-op analgesic was administered at 24 hours. Mice were given at least 2 weeks for recovery and viral expression prior to experimentation.

#### *In vivo* Ca Imaging

An upright two-photon microscope (Bruker Investigator) coupled with a Spectra Physics X3 femtosecond laser was used to acquire images at ∼29 frames/s in galvo-resonant mode. The excitation wavelength was tuned to 920 nm, and a piezoelectric motor was used to rapidly step through z axis space. The microscope was set up for in vivo imaging with a Somnosuite (isoflurane) anesthesia machine coupled to a homeothermic control warm pad (Kent Scientific) and a World Precision pump (Harvard apparatus) for infusing nutrients in the gut.

Mice fasted for at least two hours after the onset of dark were placed under continuous anesthesia (1.5% isoflurane/oxygen) and kept on a heating pad to maintain body temperature throughout the procedure. An incision was made above the sternum and below the jaw, the carotid and vagus nerve were exposed after separation of the salivary glands. Retractors were used to pull the sternomastoid, omohyoid and posterior belly of digastric muscle sideways and make the nodose ganglion visible. The vagus nerve was cut immediately above the nodose ganglion, which was carefully separated from the hypoglossal nerve and small branches. The vagus nerve was then dissected from the carotid and surrounding soft tissues, and the nodose ganglion was gently placed on a stable imaging platform consisted of a 8 mm diameter coverslip attached to a metal arm affixed to a magnetic base. Surgical silicone adhesive (Kwik-Sil, WPI) was applied onto the vagus nerve to keep it immobilized on the coverslip and the ganglion immersed in a drop of DMEM (brand) media was covered with a second coverslip. Imaging was performed) using a 20x, water immersion, upright objective.

#### Intraduodenal stimulation

Infusions were performed with a precision pump that held syringes attached to a silicone tubing and filled with either corn oil (Mazola^®^) or saline. Prior to surgery, the nodose ganglia was uncovered by making a small midline incision into the abdomen of anesthetized mice to expose the stomach. Next, a silicone tubbing was inserted through a small incision in the stomach wall, into the proximal portion of the duodenal lumen. An exit port was created ∼2cm distally to pylorus by transecting the duodenum. Super glue was applied onto the wall of the stomach to prevent the tubbing from sliding out of the intestine. Gauze soaked with saline (37°C) were placed on the stomach to keep it hydrated throughout the experiment. Recordings of baseline signals started with continuous infusion of saline for 5 minutes (100ul/min, 37°C) followed by 5 minutes of corn oil infusion (100ul/min, 37°C), and then additional 10 min of saline infusion (100ul/min) until the end of the experiment. For distension, the exit port was clamped and baseline signals were recorded for 1 minute followed by 20 seconds of duodenal inflation achieved by delivering a bolus injection of air (∼1.5ml).

#### Calcium imaging analysis

GCaMP6s fluorescent changes were outlined in regions of interests (ROIs) with each ROI defining a single cell throughout the imaging session. The pixel intensity in ROIs (average across pixels) were calculated frame by frame (ImageJ) and exported to excel for manual analysis. The baseline signal was defined as the average GCaMP6s fluorescence over 5 minutes (fat) or 1 minute (distension) period prior to the stimulus introduction. Neurons were considered responsive if 1) the peak GCaMP6s fluorescence was > 20% above the baseline mean and 2) if the increase in fluorescence is within 5 minutes (fat) or 20 second (distension) window of stimulus. Cells were considered responsive to nutrients if the following criteria were met: 1) the peak GCaMP6s fluorescence was two standard deviations above the baseline mean and 2) the mean GCaMP6s fluorescence was ≥ 20% above the baseline mean for a 20- and 3-seconds window around the peak for fat and distension, respectively. Nodose ganglia in which neurons did not present baseline activity were excluded from the study.

#### Optical fiber implantation

Mice were anesthetized mice with isoflurane and placed on a heating pad, with the head affixed to a stereotaxic apparatus (World Precision Instruments, Sarasota, FL). After skin incision (1-1.5 cm) and removal of all soft tissue from the surface of the skull, the periosteum was removed by blunt dissection. A dental drill was used to drill holes (0.61-1mm) above the target area and stainless-steel screws secured allowing better fixation of the probe. Optogenetic posts were composed of fiber optic (FT200UMT, Thorlabs, Newton, NJ) secured in a ceramic ferrule (LC Zirconia Ferrule FZI-LC-230, Kientec, Stuart, FL) with UV-cured adhesive (RapidFix, St. Louis, MO). Posts were implanted over vagal terminals in the NTS (AP: −7.5mm, ML: ± 0.3mm, DV −5.0mm) and secured with dental cement (GC Fujicem 2, GC America, Tokyo, Japan).

#### Intragastric catheter implantation

IG catheters consisted of 6 cm silicon tubing (.047” OD x .025” ID, SIL047, Braintree Scientific, Braintree, MA) with 6 beads of silicon glue (#31003, Marineland, Blacksburg, VA) applied at 1 mm, 3 mm, 13 mm, and 15 mm from the distal end, and at 10 mm and 12mm from the proximal end, and a Pinport (Instech Labs, Plymouth Meeting, PA) for chronic intermittent access.

Analgesics buprenorphine XR (1 mg/kg) and carprofen (5 mg/kg) were administered subcutaneously 20 min prior to surgery. Once animals had been anesthetized, a midline incision was made into the abdomen, and the stomach exteriorized. A purse-string suture was made at the junction of the greater curvature and fundus, avoiding major blood vessels. The center of the purse-string was punctured and the catheter was pushed in until the first drop of silicone glue was inside the stomach. Then, the purse string was tightened around the tubing, which was then tunneled subcutaneously to the dorsum via a small hole made into the abdominal muscle. The catheter was tightened and tied to the peritoneal wall between the third and fourth drops of silicone glue. A small incision to the dorsum between the shoulder plates was then made to allow for catheter exteriorization. A sterile blunt probe, 1 mm by 14 cm inserted and tunneled under the skin caudal to the left foreleg, to the abdominal incision. The catheter was threaded onto the end of the blunt probe and pulled through until the first bead on the proximal end was externalized, with the second proximal bead under the skin. Incisions were sutured and thoroughly disinfected and the exterior end of the catheter plugged. Post-op analgesic was administered at 24 hours. Mice were fed with moistened chow and given at least 1 week for recovery prior to experimentation. Daily body weight was monitored until pre-surgical weight was regained.

### Histological Procedures

#### Perfusion and tissue preparation

Mice were anesthetized with isofluorane and then perfused transcardially with phosphate buffer (PBS 0.02M, pH 7.4), followed by cold (4°C) 4% paraformaldehyde (PFA). After perfusion, tissues were harvested post-fixed in 4% PFA for 24h and kept at 4°C in a 30% sucrose in PBS solution until processing. Sterile PBS was used to perfuse tissues that were used for *in situ* hybridization.

#### Immunostaining

Tissues were embedded in OCT compound and sections (Brains, 35µm; intestines, 30µm; and nodose ganglia, 16µm) were prepared using a cryostat. Tissues slices were collected and washed 3 x 10 min with PBS, blocked (20% donkey serum, 0.1% Triton X-100 and 1% BSA in PBS) for 30 minutes, and incubated overnight at 4°C with primary antibodies (1:1000) diluted in permeabilizing solution (2% donkey serum, 0.1% Triton X-100 and 1% BSA in PBS). On the next day, tissue sections were washed 3 x 10 min with PBS, incubated for 2 hours at 37°C secondary antibodies (1:1000 in permeabilizing solution), and then washed again 3 x 10 min with PBS. Lastly, the sections were mounted on Superfrost Plus slides using antifade mountant media (Fischer, P36961) and stored in −20°C.

Antibodies used: rabbit anti-cFos (Cell Signaling; 2250), goat anti-GFP (Abcam; 305635), rabbit-anti mCherry (Takara Bio Clontech; 632496), rabbit anti-CART (55–102) (Phoenix Pharmaceuticals; H-003-62), Alexa 648-conjugated donkey anti-rabbit IgG (Abcam; ab150075) and Alexa 488-conjugated donkey anti-goat IgG (Abcam; ab150129).

#### cFos measurements

Fos immunoreactivity was used to determine the effects of intraduodenal optogenetic stimulation, nutrient infusion and/or distension on neuronal activity. Ninety minutes after the appropriate stimulation, mice were sacrificed and perfused as described earlier. Braine sections (35 µm) labelled for cFos expression were imaged using a Keyence BZ-X800. Fos expression was analyzed and quantified on coronal hindbrain sections for the presence of Fos+ neurons in the NTS, AP and DMV. Quantification of cFos expression and colocalization with tdTomato+ neurons was conducted using merged fluorescence images and analyzed using Nikon NIS Elements software.

#### RNAscope in situ hybridization

Each NDG was sectioned at 16 μm into serial sections using a Leica CM3050 S cryostat (Leica, Buffalo Grove, IL, United States). Sections were immediately mounted and slides were dipped in ethanol (100%) for a few seconds. After air-drying at room temperature for 1 h, slides were rinsed with PBS, incubated in 4% PFA for 5 min, and then dehydrated for 5 min in 50, 75, and 100% EtOH. Following that, slides are incubated in H_2_O_2_ for 10 min, rinsed with PBS, and allowed to dry for 10–15 min before being stored at –80°C for further processing.

RNAscope in situ hybridization was performed using the RNAscope® V2 Multiplex Fluorescent Reagent Kit (Advanced Cell Diagnostics, Newark, CA, United States) as per the manufacturer’s instructions. The probe CARTPT (Mm-Cartpt-C2; Cat. No. 432009-C2) was used.

#### Tissue imaging

Tissue sections of brains and NG were imaged using a Keyence BZ-X800. Image analysis of cFos expression and colocalization was conducted using Nikon NIS Elements software. For imaging of sections processed for in situ hybridization, positive and negative control probes were used to determine exposure time and image processing parameters for optimal visualization of mRNA signals and control for possible photobleaching. Whole NG samples were imaged using a 2-Photon microscope (Bruker).

#### Gut tissue clearing and quantification of vagal terminals

The duodenum (11mm) was and then opened into flat sheets with longitudinal cuts along the mesenteric attachment. The sheets of muscle wall of the duodenum were prepared as sets of wholemounts after the submucosal/mucosal layers were separated with a forceps. Tissue was cleared by incubating in TDE (2,2′-thiodiethanol, MilliporeSigma) for a minimum of 2 h prior to imaging. For imaging, cleared tissues were compressed onto the slide using a coverslip that was secured glued to it. Images were captured and processed using a laser-scanning confocal microscope (Nikon Instruments Inc., Melville, NY, United States). Multiple z stacks captured at 10 x magnification throughout the duodenum (whole-mount).

Image analysis was conducted using Nikon NIS Elements software. A counting grid was used (1mm x 1mm) and the tdTomato+ endings wrapping crypts or puncta at the base of villi were considered as mucosal innervation while IGLEs were identified using standard criteria (*1*).

#### Behavioral Studies

The following behavioral experiments were performed in mouse behavior chambers (Med Associates Inc.) equipped with two slots for sipper bottles, located on opposite sides on the same cage wall. A contact-based licking detection device was connected to each sipper and provided real-time data of licking responses (10ms resolution) that were saved in computer files for posterior analysis. To pair lick responses with intra-gastric infusions, we connected pumps equipped with TTL input devices to the behavior chambers and programmed (MED-PC IV) to automatically trigger infusions in response to the detection of licks. For optogenetic stimulation, separate chambers with a 5 - hole nose poke wall with infrared beam detectors were used for nose-poke operant tasks (Med Associates Inc).

#### Two-bottle choice Flavor-nutrient conditioning

Calorie-restricted mice (90% of body weight) were acclimated to sound proof operant lickometer boxes (MedAssociates) and trained to drink from spouts containing saccharin (0.2%), 1 h per day for five consecutive days. On the first day of flavor-preference tests, mice underwent a pre-test, in which they were given 10-min access to two artificially sweetened (0.025% w/v saccharin) Kool-Aid (0.05% w/v) flavors in mouse behavior chambers (Med Associates Inc.). At the 5-min point, the flavor positions were swapped to minimize the effects of individual side preference. The total number of licks for each flavor was computed across the session and the more preferred flavor was paired to saline, while fat (positive conditioned stimulus) was paired to the less preferred flavor the to avoid a ceiling effect on preference for the fat-paired flavor. Conditioning sessions lasted for 1 hour and were performed for 6 consecutive days, alternating daily each intragastric solution with the paired flavor. Thus, there were 3 sessions associated with each specific flavor-fat pair. After 6 days of conditioning, on post-test day, the 10-min 2-bottle preference test was repeated, exactly as during the pre-test. Outcomes were measured by the percent preference each subject had for the nutrient-paired flavor during post-test.

#### Flavor-CNO conditioning

Calorie-restricted mice (90% of body weight) were acclimated to the operant lickometer chambers (MedAssociates), trained to lick from spouts and submitted to pre-tests and post-tests as described above. Conditioning sessions lasted for 1 hour and were performed for 6 consecutive days, alternating daily i.p. injections of CNO (1mg/kg) or saline solution with the paired flavor.

#### Nose-poke task (self-stimulation)

Mice were trained to nose-poke for optogenetic stimulation in a Med Associates nose poke chamber with two slots for nose poke at symmetrical locations on one of the cage walls. Nose poke slots were connected to a photo-beam detection device, and only one of them triggered optogenetic stimulation (active side) at 1 Hz frequency for 1 s duration at 5 mW intensity (measured at the fiber tip), as previously validated (*2*). Mice were randomly assigned right or left active nose-pokes in a counterbalanced manner and underwent 3 days of 30-min nose-poke training during which a small amount of powdered rodent chow was placed in both slots to motivate mice to explore the nose pokes. In all sessions, mice were tethered to a fiber-optic cable with swivel attached to laser. Following 3 days of training, mice underwent 3 days of testing without food in the nose-pokes, during which the number of active and inactive nose pokes was measured over a 30-min test period each day.

#### Methylcellulose gavage

Adult male mice were fasted overnight before the experiment. Fasted mice then received an oral gavage of 400 µL of methyl cellulose 4000 cp (Sigma-Aldrich), diluted in water. The gavage procedure was performed using a metal feeding tube attached to a syringe, ensuring minimal stress and discomfort to the animals. Following the gavage, mice were individually placed in operant chambers (Med Associates, St. Albans, VT, USA) for a 1-hour experimental session. Each chamber was equipped with a metal sipper containing 7.5% (v/v) fat from the Microlipid emulsion (Nestle, Inc., USA) and mice were allowed to freely lick the metal sipper to consume the intralipid solution during the 1-hour test period. Licking behavior and intralipid consumption were recorded continuously throughout the 1-hour session using the Med Associates data acquisition system. The total number of licks and the volume of intralipid consumed were quantified for each mouse.

#### Food intake measurement

Meal patterns were analyzed in a BioDAQ (Research diets Inc, New Brunswick, NJ) episodic food intake monitoring system (n = 6-9/group). Meals were defined by at least 0.02 g consumed without interruption by a pause of > 5 minutes. The system consists of a low spill enlarged opening food hopper placed on an electronic balance mounted together on the animals’ cage. Chow intake was continuously measured for 2 weeks during chemogenetic tests, or 1 week in experiments with neuron ablation (Figure 4).

The animals were single housed and were habituated for one week to eating food on the BioDAQ hopper and to receiving saline injections 15-20 min before the dark onset. Drug- or neuronal manipulation-induced changes to eating behavior were assessed from dark onset in overnight fasted mice, after which food was available at ad libitum.

Clozapine N-oxide (CNO) was administered i.p. at 1 mg kg−1 for all experiments, typically 15 min before dark onset. We have previously determined that CNO at this dose does not affect eating behavior (*2*). All chemogenetic experiments in the present study were conducted in DREADD-expressing mice by using a within-subjects design in a sense that all mice received both CNO and saline in a counterbalanced manner, and hence acted as their own controls. For experiments with unilateral neuronal ablation, animals from the control group got bilateral NG injections of PHP.S-FLEX-tdTomato and AAV9-EF1a-double floxed-hChR2(H134R)-EYFP-WPRE-HGHpA in a counter-balanced manner.

#### Paracetamol absorption test (gastric emptying assay)

Mice were fasted for 4h at the onset of dark and received an i.p. injection of either CNO (1mg/kg) or saline 15 minutes before the test. Next, a solution of Ensure and Paracetamol (100 mg/kg, 250-300 ul) was administered through gavage and blood samples were obtained at 0, 15, 30, 60 and 90 minutes after gavage. Glass capillaries (WPI, cat. No. 1B100-3) coated with a drop of heparin (50U/ml) was used to collect the blood from the tip of the tail. Tubes with blood samples were then centrifuged at 1500xg for 10 minutes and plasma collected and stored in −20 C. A commercial kit (Acetaminophen L3K® Assay, Sekisui diagnostics) was used to measure the levels of paracetamol in the plasma and the samples were run in duplicates following manufacturer’s instruction.

#### Analysis of whole-nodose scSeq data

We used the raw counts of the single-cell RNA sequencing from the left and right nodose ganglia previously published (*3*) (GEO accession number GSE185173). All the analysis were performed using the Seurat version 4.0 (Hao et al., 2021) for R (version 4.1.3) and RStudio (2022.02.1 Build 461). Data normalization, transformation, scaling, linear dimensional reduction and cell clustering were performed using default functions provided in Seurat. The predicted cell identities published in (*3*) and gene markers of nodose ganglion cells published in (*4*) were used as a reference to subset the cells of interest in downstream analysis.

#### Slice electrophysiology

Mice were deeply anesthetized with 4% isoflurane, followed by abdominal hair removal and sterilization. The animals were then positioned supine on a warming pad, and a minor midline incision was created to expose both the stomach and duodenum. Subsequently, silicone tubing was inserted via a small opening in the stomach wall into the proximal section of the duodenal lumen. Throughout the experiment, the gut was maintained hydrated by applying saline-soaked gauze (37°C). To induce duodenal distension, a 0.5 ml bolus air injection was administered. For nutrient stimulation, the duodenum received a 1-minute infusion of 50µl corn oil. Post-stimulation, incisions were sutured, and the mice were allowed to recover on a heating pad until they voluntarily moved to the unheated section of the cage. After 30 min, the brain was collected, glued onto the cutting stage and submersed in ice-cold, oxygenated aCSF (equilibrated with 95% O2-5% CO2). Coronal or horizontal brain slices (200 μm) containing the substantia nigra compacta were cut using a MicroSlicer Zero 1N (Dosaka, Kyoto, Japan).

Spontaneous firing activity of midbrain dopamine neurons was examined via whole cell current clamp recordings as previously described (*5–7*). The neurons were continuously perfused with artificial cerebral spinal fluid (aCSF) containing (in mM): 126 NaCl, 2.5 KCl, 2 CaCl2, 26 NaHCO3, 1.25 NaH2PO4, 2 MgSO4, and 10 dextrose, equilibrated with 95% O2-5% CO2; pH was adjusted to 7.4 at 37°C. Patch electrodes were fabricated from borosilicate glass (1.5 mm outer diameter; World Precision Instruments, Sarasota, FL) with the P-2000 puller (Sutter Instruments, Novato, CA). The tip resistance was in the range of 3-5 MΩ. The electrodes were filled with a pipette solution containing (in mM): 120 potassium-gluconate, 20 KCl, 2 MgCl2, 10 HEPES, 0.1 EGTA, 2 ATP, and 0.25 GTP, with pH adjusted to 7.25 with KOH. All experiments were performed at 37°C. To standardize action potential (AP) recordings, neurons were held at their resting membrane potential (see below) by DC application through the recording electrode. Action potential was recorded if the following criteria were met: a resting membrane potential polarized than −35 mV and an action potential peak amplitude of >60 mV. Action potential was measured using Clampfit 10 software (Axon instruments, Foster City, CA). Steady-state basal activity was recorded for 2–3 min before bath application of the drug. The spontaneous spike activity of dopamine neurons was obtained by averaging 1 min interval activities at baseline (before manganese) and after 3-5 min of drugs.

The electrophysiology data were acquired using the ClampEx 10 software (Molecular Devices). The data were analyzed offline using pClamp 10. For all experiments

#### Data Analysis

Statistical analysis for the experiments is described in each figure legend and was determined using GraphPad Prism 9 software. Two-tailed unpaired Student’s t tests were used for comparing two groups; Two-tailed paired Student’s t test was used for comparing two treatments or tests in the same animal. One-way ANOVA, with or without repeated-measures, was used for comparing three groups; two-way ANOVA, with or without repeated-measures, was used for comparing more than one factor between. Data are presented as mean ± SEM and statistical significance is declared at p < 0.05.

## REFERENCES

1. H.-R. Berthoud, The vagus nerve, food intake and obesity. Regul Pept. 149, 15–25 (2008).

2. G. J. Dockray, G. Burdyga, Plasticity in vagal afferent neurones during feeding and fasting: mechanisms and significance. Acta Physiol (Oxf). 201, 313–321 (2011).

3. H. R. Berthoud, M. Kressel, H. E. Raybould, W. L. Neuhuber, Vagal sensors in the rat duodenal mucosa: distribution and structure as revealed by in vivo DiI-tracing. Anat Embryol (Berl). 191, 203–212 (1995).

4. H. R. Berthoud, L. M. Patterson, F. Neumann, W. L. Neuhuber, Distribution and structure of vagal afferent intraganglionic laminar endings (IGLEs) in the rat gastrointestinal tract. Anat Embryol (Berl). 195, 183–191 (1997).

5. F. B. Wang, T. L. Powley, Topographic inventories of vagal afferents in gastrointestinal muscle. J Comp Neurol. 421, 302–324 (2000).

6. T. L. Powley, R. A. Spaulding, S. A. Haglof, Vagal afferent innervation of the proximal gastrointestinal tract mucosa: chemoreceptor and mechanoreceptor architecture. J Comp Neurol. 519, 644–660 (2011).

7. Q. Zhao, C. D. Yu, R. Wang, Q. J. Xu, R. Dai Pra, L. Zhang, R. B. Chang, A multidimensional coding architecture of the vagal interoceptive system. Nature. 603, 878–884 (2022).

8. S. L. Prescott, B. D. Umans, E. K. Williams, R. D. Brust, S. D. Liberles, An Airway Protection Program Revealed by Sweeping Genetic Control of Vagal Afferents. Cell. 181, 574–589.e14 (2020).

9. L. Bai, S. Mesgarzadeh, K. S. Ramesh, E. L. Huey, Y. Liu, L. A. Gray, T. J. Aitken, Y. Chen, L. R. Beutler, J. S. Ahn, L. Madisen, H. Zeng, M. A. Krasnow, Z. A. Knight, Genetic Identification of Vagal Sensory Neurons That Control Feeding. Cell. 179, 1129–1143.e23 (2019).

10. J. Kupari, M. Häring, E. Agirre, G. Castelo-Branco, P. Ernfors, An Atlas of Vagal Sensory Neurons and Their Molecular Specialization. Cell Rep. 27, 2508–2523.e4 (2019).

11. H. Suzuki, T. R. Thiele, S. Faumont, M. Ezcurra, S. R. Lockery, W. R. Schafer, Functional asymmetry in Caenorhabditis elegans taste neurons and its computational role in chemotaxis. Nature. 454, 114–117 (2008).

12. O. Güntürkün, F. Ströckens, S. Ocklenburg, Brain Lateralization: A Comparative Perspective. Physiol Rev. 100, 1019–1063 (2020).

13. K. L. Buchanan, L. E. Rupprecht, M. M. Kaelberer, A. Sahasrabudhe, M. E. Klein, J. A. Villalobos, W. W. Liu, A. Yang, J. Gelman, S. Park, P. Anikeeva, D. V. Bohórquez, The preference for sugar over sweetener depends on a gut sensor cell. Nat Neurosci. 25, 191–200 (2022).

14. C. Broberger, K. Holmberg, M. J. Kuhar, T. Hökfelt, Cocaine- and amphetamine-regulated transcript in the rat vagus nerve: A putative mediator of cholecystokinin-induced satiety. Proc Natl Acad Sci U S A. 96, 13506–13511 (1999).

15. S. J. Lee, J.-P. Krieger, M. Vergara, D. Quinn, M. McDougle, A. de Araujo, R. Darling, B. Zollinger, S. Anderson, A. Pan, E. J. Simonnet, A. Pignalosa, M. Arnold, A. Singh, W. Langhans, H. E. Raybould, G. de Lartigue, Blunted Vagal Cocaine- and Amphetamine-Regulated Transcript Promotes Hyperphagia and Weight Gain. Cell Rep. 30, 2028–2039.e4 (2020).

16. A. Singh, A. M. de Araujo, J.-P. Krieger, M. Vergara, C. K. Ip, G. de Lartigue, Demystifying functional role of cocaine- and amphetamine-related transcript (CART) peptide in control of energy homeostasis: A twenty-five year expedition. Peptides. 140, 170534 (2021).

17. H. R. Berthoud, W. L. Neuhuber, Functional and chemical anatomy of the afferent vagal system. Auton Neurosci. 85, 1–17 (2000).

18. V. P. Zagorodnyuk, B. N. Chen, S. J. Brookes, Intraganglionic laminar endings are mechano-transduction sites of vagal tension receptors in the guinea-pig stomach. J Physiol. 534, 255–268 (2001).

19. E. K. Williams, R. B. Chang, D. E. Strochlic, B. D. Umans, B. B. Lowell, S. D. Liberles, Sensory Neurons that Detect Stretch and Nutrients in the Digestive System. Cell. 166, 209–221 (2016).

20. M. McDougle, A. de Araujo, M. Vergara, M. Yang, A. Singh, I. Braga, N. Urs, B. Warren, G. de Lartigue, Labeled lines for fat and sugar reward combine to promote overeating (2022), p. 2022.08.09.503218,, doi:10.1101/2022.08.09.503218.

21. A. Sclafani, Conditioned food preferences. Bull. Psychon. Soc. 29, 256–260 (1991).

22. S. E. Thanarajah, H. Backes, A. G. DiFeliceantonio, K. Albus, A. L. Cremer, R. Hanssen, R. N. Lippert, O. A. Cornely, D. M. Small, J. C. Brüning, M. Tittgemeyer, Food Intake Recruits Orosensory and Post-ingestive Dopaminergic Circuits to Affect Eating Desire in Humans. Cell Metab. 29, 695–706.e4 (2019).

23. W. Han, L. A. Tellez, M. H. Perkins, I. O. Perez, T. Qu, J. Ferreira, T. L. Ferreira, D. Quinn, Z.-W. Liu, X.-B. Gao, M. M. Kaelberer, D. V. Bohórquez, S. J. Shammah-Lagnado, G. de Lartigue, I. E. de Araujo, A Neural Circuit for Gut-Induced Reward. Cell. 175, 665–678.e23 (2018).

24. G. de Lartigue, Role of the vagus nerve in the development and treatment of diet-induced obesity. J Physiol. 594, 5791–5815 (2016).

25. C. J. Guenthner, K. Miyamichi, H. H. Yang, H. C. Heller, L. Luo, Permanent genetic access to transiently active neurons via TRAP: targeted recombination in active populations. Neuron. 78, 773–784 (2013).

26. A. Geliebter, S. Westreich, S. A. Hashim, D. Gage, Gastric balloon reduces food intake and body weight in obese rats. Physiology & Behavior. 39, 399–402 (1987).

27. R. J. Phillips, T. L. Powley, Gastric volume rather than nutrient content inhibits food intake. Am J Physiol. 271, R766–769 (1996).

28. H. R. Berthoud, T. L. Powley, Vagal afferent innervation of the rat fundic stomach: morphological characterization of the gastric tension receptor. J Comp Neurol. 319, 261–276 (1992).

29. T. Ichiki, T. Wang, A. Kennedy, A.-H. Pool, H. Ebisu, D. J. Anderson, Y. Oka, Sensory representation and detection mechanisms of gut osmolality change. Nature. 602, 468–474 (2022).

30. A. Iggo, Tension receptors in the stomach and the urinary bladder. J Physiol. 128, 593–607 (1955).

31. D. I. Brierley, M. K. Holt, A. Singh, A. de Araujo, M. McDougle, M. Vergara, M. H. Afaghani, S. J. Lee, K. Scott, C. Maske, W. Langhans, E. Krause, A. de Kloet, F. M. Gribble, F. Reimann, L. Rinaman, G. de Lartigue, S. Trapp, Central and peripheral GLP-1 systems independently suppress eating. Nat Metab. 3, 258–273 (2021).

32. C. Ran, J. C. Boettcher, J. A. Kaye, C. E. Gallori, S. D. Liberles, A brainstem map for visceral sensations. Nature. 609, 320–326 (2022).

33. D.-Y. Kim, G. Heo, M. Kim, H. Kim, J. A. Jin, H.-K. Kim, S. Jung, M. An, B. H. Ahn, J. H. Park, H.-E. Park, M. Lee, J. W. Lee, G. J. Schwartz, S.-Y. Kim, A neural circuit mechanism for mechanosensory feedback control of ingestion. Nature. 580, 376–380 (2020).

34. M. F. Gonzalez, J. A. Deutsch, Vagotomy abolishes cues of satiety produced by gastric distension. Science. 212, 1283–1284 (1981).

35. R. J. Phillips, T. L. Powley, Gastric volume detection after selective vagotomies in rats. Am J Physiol. 274, R1626–1638 (1998).

36. D. Borgmann, E. Ciglieri, N. Biglari, C. Brandt, A. L. Cremer, H. Backes, M. Tittgemeyer, F. T. Wunderlich, J. C. Brüning, H. Fenselau, Gut-brain communication by distinct sensory neurons differently controls feeding and glucose metabolism. Cell Metab. 33, 1466–1482.e7 (2021).

37. T. Zhang, M. H. Perkins, H. Chang, W. Han, I. E. de Araujo, An inter-organ neural circuit for appetite suppression. Cell. 185, 2478–2494.e28 (2022).

38. R. A. Travagli, G. E. Hermann, K. N. Browning, R. C. Rogers, Musings on the wanderer: what’s new in our understanding of vago-vagal reflexes? III. Activity-dependent plasticity in vago-vagal reflexes controlling the stomach. Am J Physiol Gastrointest Liver Physiol. 284, G180–187 (2003).

39. M. B. Parent, S. Higgs, L. G. Cheke, S. E. Kanoski, Memory and eating: A bidirectional relationship implicated in obesity. Neurosci Biobehav Rev. 132, 110–129 (2022).

40. A. N. Suarez, T. M. Hsu, C. M. Liu, E. E. Noble, A. M. Cortella, E. M. Nakamoto, J. D. Hahn, G. de Lartigue, S. E. Kanoski, Gut vagal sensory signaling regulates hippocampus function through multi-order pathways. Nat Commun. 9, 2181 (2018).

41. H. Cai, W. Haubensak, T. E. Anthony, D. J. Anderson, Central amygdala PKC-δ(+) neurons mediate the influence of multiple anorexigenic signals. Nat Neurosci. 17, 1240–1248 (2014).

42. J. A. Hardaway, L. R. Halladay, C. M. Mazzone, D. Pati, D. W. Bloodgood, M. Kim, J. Jensen, J. F. DiBerto, K. M. Boyt, A. Shiddapur, A. Erfani, O. J. Hon, S. Neira, C. M. Stanhope, J. A. Sugam, M. P. Saddoris, G. Tipton, Z. McElligott, T. C. Jhou, G. D. Stuber, M. R. Bruchas, C. M. Bulik, A. Holmes, T. L. Kash, Central Amygdala Prepronociceptin-Expressing Neurons Mediate Palatable Food Consumption and Reward. Neuron. 102, 1088 (2019).

43. M. J. F. Robinson, S. M. Warlow, K. C. Berridge, Optogenetic excitation of central amygdala amplifies and narrows incentive motivation to pursue one reward above another. J Neurosci. 34, 16567– 16580 (2014).

44. D. Burdakov, D. Peleg-Raibstein, The hypothalamus as a primary coordinator of memory updating. Physiol Behav. 223, 112988 (2020).

45. G. D. Stuber, R. A. Wise, Lateral hypothalamic circuits for feeding and reward. Nat Neurosci. 19, 198–205 (2016).

46. H. D. Critchley, The human cortex responds to an interoceptive challenge. Proceedings of the National Academy of Sciences. 101, 6333–6334 (2004).

47. N. Medford, H. D. Critchley, Conjoint activity of anterior insular and anterior cingulate cortex: awareness and response. Brain Struct Funct. 214, 535–549 (2010).

48. A. Bisazza, C. Cantalupo, M. Capocchiano, G. Vallortigara, Population lateralisation and social behaviour: a study with 16 species of fish. Laterality. 5, 269–284 (2000).

49. L. J. Rogers, P. Zucca, G. Vallortigara, Advantages of having a lateralized brain. Proc Biol Sci. 271 **Suppl 6**, S420–422 (2004).

50. T. E. Zandstra, R. G. E. Notenboom, J. Wink, P. Kiès, H. W. Vliegen, A. D. Egorova, M. J. Schalij, M. C. De Ruiter, M. R. M. Jongbloed, Asymmetry and Heterogeneity: Part and Parcel in Cardiac Autonomic Innervation and Function. Front Physiol. 12, 665298 (2021).

51. H.-R. Berthoud, Anatomy and function of sensory hepatic nerves. Anat Rec A Discov Mol Cell Evol Biol. 280, 827–835 (2004).

52. S. B. Mazzone, B. J. Undem, Vagal Afferent Innervation of the Airways in Health and Disease. Physiol Rev. 96, 975–1024 (2016).

53. K. E. Fasanella, J. A. Christianson, R. S. Chanthaphavong, B. M. Davis, Distribution and Neurochemical Identification of Pancreatic Afferents in the Mouse. J Comp Neurol. 509, 42–52 (2008).

54. N. Jayaprakash, W. Song, V. Toth, A. Vardhan, T. Levy, J. Tomaio, K. Qanud, I. Mughrabi, Y.-C. Chang, M. Rob, A. Daytz, A. Abbas, Z. Nassrallah, B. T. Volpe, K. J. Tracey, Y. Al-Abed, T. Datta-Chaudhuri, L. Miller, M. F. Barbe, S. C. Lee, T. P. Zanos, S. Zanos, Organ- and function-specific anatomical organization of vagal fibers supports fascicular vagus nerve stimulation. Brain Stimul. 16, 484–506 (2023).

55. N. Thompson, S. Mastitskaya, D. Holder, Avoiding off-target effects in electrical stimulation of the cervical vagus nerve: Neuroanatomical tracing techniques to study fascicular anatomy of the vagus nerve. J Neurosci Methods. 325, 108325 (2019).

56. N. Thompson, E. Ravagli, S. Mastitskaya, F. Iacoviello, T.-R. Stathopoulou, J. Perkins, P. R. Shearing, K. Aristovich, D. Holder, Organotopic Organization of the Cervical Vagus Nerve (2023), p. 2022.02.24.481810,, doi:10.1101/2022.02.24.481810.

57. A. R. Upadhye, C. Kolluru, L. Druschel, L. Al Lababidi, S. S. Ahmad, D. M. Menendez, O. N. Buyukcelik, M. L. Settell, S. L. Blanz, M. W. Jenkins, D. L. Wilson, J. Zhang, C. Tatsuoka, W. M. Grill, N. A. Pelot, K. A. Ludwig, K. J. Gustafson, A. J. Shoffstall, Fascicles split or merge every ∼560 microns within the human cervical vagus nerve. J Neural Eng. 19 (2022), doi:10.1088/1741-2552/ac9643.

58. A. N. Pouliopoulos, M. F. Murillo, R. L. Noel, A. J. Batts, R. Ji, N. Kwon, H. Yu, C.-K. Tong, J. N. Gelinas, D. K. Araghy, S. A. Hussaini, E. E. Konofagou, Non-invasive optogenetics with ultrasound-mediated gene delivery and red-light excitation. Brain Stimul. 15, 927–941 (2022).

59. B. Hsueh, R. Chen, Y. Jo, D. Tang, M. Raffiee, Y. S. Kim, M. Inoue, S. Randles, C. Ramakrishnan, S. Patel, D. K. Kim, T. X. Liu, S. H. Kim, L. Tan, L. Mortazavi, A. Cordero, J. Shi, M. Zhao, T. T. Ho, A. Crow, A.-C. W. Yoo, C. Raja, K. Evans, D. Bernstein, M. Zeineh, M. Goubran, K. Deisseroth, Cardiogenic control of affective behavioural state. Nature. 615, 292–299 (2023).

## REFERENCES

1. F. B. Wang, T. L. Powley, Topographic inventories of vagal afferents in gastrointestinal muscle. J Comp Neurol. 421, 302–324 (2000).

2. W. Han, L. A. Tellez, M. H. Perkins, I. O. Perez, T. Qu, J. Ferreira, T. L. Ferreira, D. Quinn, Z.-W. Liu, X.-B. Gao, M. M. Kaelberer, D. V. Bohórquez, S. J. Shammah-Lagnado, G. de Lartigue, I. E. de Araujo, A Neural Circuit for Gut-Induced Reward. Cell. 175, 665–678.e23 (2018).

3. K. L. Buchanan, L. E. Rupprecht, M. M. Kaelberer, A. Sahasrabudhe, M. E. Klein, J. A. Villalobos, W. W. Liu, A. Yang, J. Gelman, S. Park, P. Anikeeva, D. V. Bohórquez, The preference for sugar over sweetener depends on a gut sensor cell. Nat Neurosci. 25, 191–200 (2022).

4. J. Kupari, M. Häring, E. Agirre, G. Castelo-Branco, P. Ernfors, An Atlas of Vagal Sensory Neurons and Their Molecular Specialization. Cell Rep. 27, 2508–2523.e4 (2019).

5. M. Lin, D. Sambo, H. Khoshbouei, Methamphetamine Regulation of Firing Activity of Dopamine Neurons. J Neurosci. 36, 10376–10391 (2016).

6. K. Saha, D. Sambo, B. D. Richardson, L. M. Lin, B. Butler, L. Villarroel, H. Khoshbouei, Intracellular methamphetamine prevents the dopamine-induced enhancement of neuronal firing. J Biol Chem. 289, 22246–22257 (2014).

7. D. O. Sambo, M. Lin, A. Owens, J. J. Lebowitz, B. Richardson, D. A. Jagnarine, M. Shetty, M. Rodriquez, T. Alonge, M. Ali, J. Katz, L. Yan, M. Febo, L. K. Henry, A. W. Bruijnzeel, L. Daws, H. Khoshbouei, The sigma-1 receptor modulates methamphetamine dysregulation of dopamine neurotransmission. Nat Commun. 8, 2228 (2017).

